# Improving drug discovery using image-based multiparametric analysis of the epigenetic landscape

**DOI:** 10.1101/541151

**Authors:** Chen Farhy, Santosh Hariharan, Jarkko Ylanko, Luis Orozco, Fu-Yue Zeng, Ian Pass, Fernando Ugarte, Camilla Forsberg, Chun-Teng Huang, David Andrews, Alexey V. Terskikh

## Abstract

With the advent of automatic cell imaging and machine learning, high-content phenotypic screening has become the approach of choice for drug discovery because it can extract drug-specific multi-layered data, which could be compared to known profiles. In the field of epigenetics, such screening approaches have suffered from a lack of tools sensitive to selective epigenetic perturbations. Here we describe a novel approach, Microscopic Imaging of Epigenetic Landscapes (MIEL), which captures the nuclear staining patterns of epigenetic marks (e.g., acetylated and methylated histones) and employs machine learning to accurately distinguish between such patterns. We validated the fidelity and robustness of the MIEL platform across multiple cells lines using dose-response curves. We employed MIEL to uncover the mechanism by which bromodomain inhibitors synergize with temozolomide-mediated killing of human glioblastoma lines. To explore alternative, non-cytotoxic, glioblastoma treatment, we screen the Prestwick chemical library and documented the power of MIEL platform to identify epigenetically active drugs and accurately rank them according to their ability to produce epigenetic and transcriptional alterations consistent with the induction of glioblastoma differentiation.

## Introduction

A cell’s epigenetic landscape is largely determined by its chromatin organization, the pattern of its DNA, and its histone modifications, all of which confer differential accessibility to areas of the genome and, through direct and indirect regulation of all DNA-related processes, form the basis of the cellular phenotype (1, 2). By collecting global information about the epigenetic landscape, for example using ATAC- or histone ChIP-seq, we can derive multilayered information regarding cellular states (3, 4). These include stable cell phenotypes such as quiescence, senescence, or cell fate, as well as transient changes such as those induced by cytokines and chemical compounds. However, current methods for collecting such information are not adapted for high-content drug screening. Over the past decade the decreasing cost and remarkable scalability have made high content screening particularly attractive for drug discovery. More recently, novel image analysis coupled with multiparametric analysis and machine learning have significantly impacted our ability to understand and process phenotypic screening outputs (5, 6). Despite these advantages, such assays have not been adapted to extract and utilize information for the cellular epigenetic landscape.

While malignant glioblastoma is the most common and lethal brain tumor, current therapeutic options offer little prognostic improvement, so the median survival time has remained virtually unchanged for decades (7–9). Tumor-propagating cells (TPCs) are a subpopulation of glioblastoma cells with increased tumorigenic capability (10) and are operationally defined as early-passaged (<15) glioblastoma cells propagated in serum-free medium (11). Compared to the bulk of the tumor, TPCs are more resistant to drugs, such as temozolomide (TMZ) and radiation therapy (12, 13). This resistance may explain the failure of traditional therapeutic strategies based on cytotoxic drugs targeting glioblastoma. Multiple approaches aimed at reducing or circumventing the resilience of TPCs have been proposed. These include targeting epigenetic enzymes (i.e., enzymes that write, remove, or read DNA and histone modifications) that would increase sensitivity to cytotoxic treatments (14–17), differentiating TPCs to reduce tumor expansion by decreased cell proliferation, and increasing sensitivity to cytotoxic treatments (18–23).

Culturing primary GBM cells in serum-containing medium induces their differentiation into cells with drastically reduced tumorigenic potential (24). In addition, Bone Morphogenetic Protein 4 (BMP4) treatment was reported to induce GBM differentiation (25, 26), which might be reversible (27) and is contingent on the presence of functional BMP receptors (28). These observations support the potential therapeutic value of small molecules that mimic the differentiation effect of serum and BMPs on TPCs

Here we have used the novel high-content screening platform MIEL to profile chromatin organization by using the endogenous patterns of histone modifications present in all eukaryotic cells. We validated this platform across multipole cell lines using epigenetically active compounds and applied MIEL to realize the mechanism by which BET inhibitors synergize with TMZ treatment. Also relevant to the tumor differentiation paradigm, we demonstrated that MIEL can identify epigenetically active drugs, classify them by molecular function, and accurately rank them by the ability to produce a set of desired epigenetic alterations consistent with inducing glioblastoma differentiation.

## Results

### Development of the MIEL platform

We have developed a novel phenotypic screening platform, MIEL, which interrogates the epigenetic landscape at both population and single cell levels using image derived features and machine learning (29). MIEL takes advantage of epigenetic marks such as histone methylation and acetylation, which are always present in eukaryotic nuclei and can be revealed by immunostaining. MIEL analyzes the immunolabeling patterns of epigenetic marks using conventional image analysis methods for nuclei segmentation, feature extraction, and previously described machine-learning algorithms (30) (Fig. 1a and Methods). Primarily, we utilized four histone modifications: H3K27me3 and H3K9me3, which are associated with condensed (closed) facultative and constitutive heterochromatin, respectively; H3K27ac, associated with transcriptionally active (open) areas of chromatin, especially at promoter and enhancer regions; and H3K4me1, associated with enhancers and other chromatin regions (Fig. 1a; (31, 32)). To focus on the intrinsic pattern of epigenetic marks, we used only texture-associated features (e.g., Haralick’s texture features (33), threshold adjacency statistics, and radial features (34)) for multivariate analysis. Previous studies have successfully employed similar features for cell painting techniques combined with multivariate analyses to accurately classify subcellular localization of proteins (34), cellular subpopulations (35), and drug mechanisms of action (30, 36-38).

**Fig. 1:**
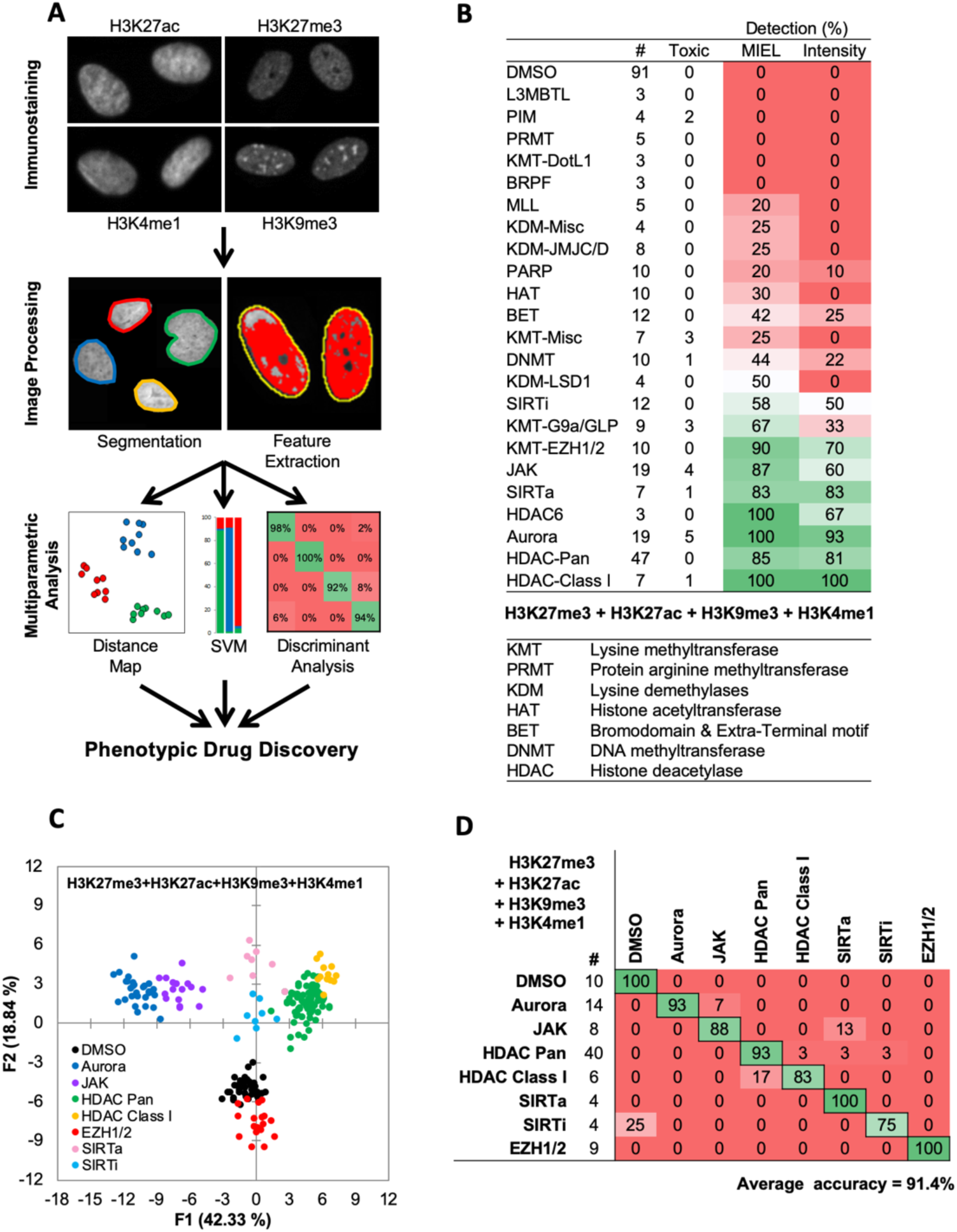
MIEL compares the epigenetic landscape of multiple cell populations and can be used to detect active epigenetic drugs across cell lines and drug concentrations. (A) Flowchart of MIEL pipeline. Fixed cells were immunostained for the desired epigenetic modifications and imaged. Nuclei were segmented based on DNA staining (Hoechst 33342 or DAPI) and texture features were calculated from the pattern of immunofluorescence. The relative similarity of multiple cell populations was assessed by calculating the multi-parametric Euclidean distance between populations centers, and represented in 2D following MDS (distance map). Discriminant analysis is used to functionally classify whole cell populations based on their multi-parametric centers. SVM classification is used to separate single cells in each population and estimate populations overlap. (B) Table showing the fraction of epigenetic drugs in each functional category identified as active by either MIEL analysis employing texture features derived from images of GBM2 cells stained for H3K9me3, H3K4me1, H3K27ac, H3K27me3, or by intensity-based analysis using the same modifications (see Methods). (C,D) Quadratic discriminant analysis using texture features derived from images of GBM2 cells treated with either DMSO or 85 active compounds (2 technical replicates per compound; 38 DMSO replicates) stained for H3K9me3, H3K27me3, H3K4me1, H3K27ac. (C) Scatter plots depicting the first 2 discriminant factors derived from features of all four histone modification images for each cell population. (D) Confusion matrix showing classification results of discriminant analysis. Left column details number of compounds or DMSO replicates for each category in the test set (1 replicate per compound). Numbers represent the percent of compounds classified correctly (diagonal) and incorrectly (off the diagonal).

We employed three main methods of data visualization and analysis: To visualize similarity between multiple cell populations, we calculated the multivariate centroids for each cell population and the Euclidean distance between all populations. To reduce data dimensionality and present as a 2D scatter plot (termed distance map), we used multidimensional scaling (MDS; Methods and Fig. 1a). To classify multiple cell populations, we employed quadratic discriminant analysis of multivariate centroids, while single cells across cell populations were classified using a Support Vector Machine (SVM; Methods and Fig. 1a).

The most commonly used cellular assays for epigenetic drug discovery are lysis and ELISA, such as AlphaLISA (PerkinElmer). Imaging-based alternatives rely on staining for relevant histone modification and monitoring changes in average fluorescent intensity (39, 40). Using MIEL, we screened a library of 222 epigenetically active compounds, many with known targets among epigenetic writers, erasers, or readers (SBP epigenetic library, Supplementary Fig. 1a, b). We focused on MIEL’s ability to (1) detect active compounds; (2) group drugs by function and identify off-target effects; (3) be robust across cell lines and drug concentrations; (4) rank active drugs, and derive information regarding drug mechanism of action.

### MIEL improves detection of epigenetically active drugs

To determine how well MIEL could detect active compounds and compare them against other intensity-based methods, primary-derived TPCs (GBM2 cell line) were treated with epigenetically active drugs for 24 hours (10 uM, triplicates). Treated cells were immunolabeled for multiple histone modifications expected to exhibit alterations following drug treatment (H3K9me3, H3K27me3, H3K27ac, and H3K4me1). Image analysis, including nuclei segmentation and features extraction, was conducted, as previously described (30) on an Acapella 2.6 (PerkinElmer). Phenotypic profiles were generated for each compound or control-treated (DMSO) treated wells. These are vectors were composed of 1048 (262 features per modification X 4 modifications) texture features derived from the staining of each modified histone modification and representing the average value for each feature across all stained cells in each cell population (drug or DMSO). When treatment reduced cell count to under 50 imaged nuclei per well, the compound was deemed toxic and excluded from analysis. Following feature normalization by z-score, we calculated the Euclidean distance between vectors of the compounds and DMSO-treated cells. These distances were then normalized (z-score) to the average distance between DMSO replicates and the standard deviation of these distances. Compounds with a distance z-score of greater than 3 were defined as active (see Methods section). This analysis identified 122 compounds that induced significant epigenetic changes. Active compounds were not uniformly distributed across all functional drug categories. Rather, we identified 10 categories in which 50% of the drugs were identified as active and nontoxic and 13 categories in which 25% or less fewer of the drugs induced detectable epigenetic alterations following a 24-hour treatment (Fig. 1b).

To compare MIEL with current thresholding methods, we repeated the calculation using mean fluorescence intensity for all histone modifications by normalizing (z-score) each drug against DMSO; active compounds were defined as compounds for which z-scored intensity for at least one of the histone modifications was greater than 3 or less than −3. As a result, we identified 94 active compounds, which were distributed across functional categories similarly to MIEL-identified compounds (Fig. 1b). For each functional category, the number of compounds identified as active using thresholding was fewer than the number identified using MIEL (Fig. 1b), demonstrating MIEL’s increased detection sensitivity over standard thresholding.

To determine the contribution of individual histone modifications, we repeated both MIEL and thresholding analyses individually for each of the 4 modifications. Using MIEL-based analysis, a single modification yielded similar detection rates to the combination of modifications across most functional categories (Supplementary Fig. 2a). Using intensity-based analysis, individual modifications yielded lower detection rates compared to the combination of modifications and displayed equal or reduced detection rates when compared to MIEL in all categories and modifications (Supplementary Fig. 2a). Of note, 3 of the 4 modifications used for MIEL analysis showed similar detection rates across most of the functional categories. However, detection rates of modified H3K27me3 were consistently reduced across the most active categories (Supplementary Fig. 2a) except for EZH1/2 inhibitors, possibly due to the role these enzymes play in regulating this posttranslational modification. To further compare MIEL and thresholding, we estimated the magnitude of epigenetic alterations induced by active compounds. We calculated the fold increase in distance from DMSO (normalized to average distance between DMSO replicates), as well as the fold change in fluorescence intensity for active compounds in each category. In all categories, MIEL showed an increased effect (Supplementary Fig. 2b).

These results demonstrate that, across all tested epigenetic modifications, detecting epigenetically active compounds using high content imaging was markedly improved by implementing MIEL compared to current image-based thresholding methods.

### MIEL suggests functional groups and identifies the off-target effects

One key advantage of phenotypic profiling methods like MIEL is the ability to classify compounds by function and identify its nonspecific effects by comparing with profiles of well-defined controls. To assess whether MIEL could correctly group compounds by function, we applied discriminant analysis (DA) to all active, nontoxic compounds from categories that had at least 3 such compounds (85 compounds; 7 categories and DMSO). Two replicates from each drug and 38 DMSO replicates were used as a training set for a quadratic DA, using all texture features derived from images of the four histone modifications (features displaying multicollinearity were reduced). The third replicate for each compound, as well as 10 DMSO replicates, were used as a test set to validate the model. Results showed that MIEL separated multiple categories of epigenetically active drugs with an average accuracy of 91.4% (Fig. 1c, d). Although many of the epigenetically active compounds induced alterations in average fluorescence (Supplementary Fig. 2b), a DA utilizing intensity measurements from all 4 channels was ineffective at separating the various categories and yielded only 51.6% average accuracy (Supplementary Fig. 3a). To test whether modification textures of individual histones contained sufficient information to distinguish between the various drug classes, we performed DA using features derived from each modification. Although this degraded MIEL’s ability to separate compound subclasses, which affected similar changes in histone modification such as Class I and Pan HDAC inhibitors, MIEL was still able to separate major categories, such as histone phosphorylation and deacetylation (Supplementary Fig. 3b).

Of note, the compound library used in this study included Pan HDAC inhibitors (HDACi), Class I HDACi, and Class I HDACi, known to also target HDAC6. HDAC inhibitors targeting both Class I and HDAC 6 displayed the same profile as Pan HDAC, and DA showed the two categories to be undistinguishable. This was likely due to the high expression of HDAC Class I and HDAC 6 and low expression of other HDACs in GBM2 glioblastoma line (Supplementary Fig. 4a, b, c).

Of the 85 compounds tested, 7 (8.2%) were identified as active but were misclassified by MIEL. One of these was valproic acid, a commonly used anticonvulsant (41), which also functions as a Pan HDAC inhibitor at high concentrations (42). Though valproic acid is expected to inhibit HDACs only at high concentrations (>1.2mM), a short 24-hour treatment induced detectable epigenetic changes even at low concentrations (<30uM). However, when we quantified H3K27ac and H3K27me3 immunofluorescence intensity at these concentrations, no increase in histone acetylation or decrease in histone methylation similar to other Pan HDAC inhibitors (TSA, SAHA; Supplementary Fig. 5a) was seen. To test, whether observed epigenetic changes resulted in corresponding transcriptomic alterations, we sequenced RNA from GBM2 cells treated with either DMSO, TSA, SAHA or valproic acid (15µM) for 24 hours and identified all genes altered by at least one of the drugs (as compared to DMSO; 118 genes). The Pan HDAC inhibitors induced similar transcriptomic changes; these were not reflected in the transcriptomic profile of valproic acid-treated cells (Supplementary Fig. 5b). To test whether MIEL profiles reflected global drug-induced transcriptomic profiles, FPKM values for all expressed genes (FPKM>1 in at least one cell population) were used to calculate the Euclidean distance between all 4 cell populations. FPKM-based distances were then correlated to image texture feature-based distances, which yielded a high and significant correlation between these metrics (R=0.91, pv<0.05; Supplementary Fig. 5c).

Taken together, we have demonstrated a unique ability of the MIEL approach to identify epigenetically active compounds, to accurately categorize them according to their molecular mechanism of action, and to detect off-target effects of compounds with known mechanism of action.

### Unbiased detection of drug concentration effect on cellular epigenetic state

As drugs vary in potency, predicting the function of unknown drugs relies on generating functional category-specific profiles that remain valid over a range of activity levels. To determine whether MIEL could correctly identify the functional category of drugs with different potencies, we treated GBM2 cells with drugs from several active categories at a range of concentrations (0.1, 0.3, 1, 3, 10µM) and conducted DA aimed at separating the different concentrations in each class. We found that for most drug categories (inhibitors of: Aurora, JAK, SIRT and EZH1/2), DA yielded low-average accuracy (Fig. 2a - Aurora kinase: 43.3%; Supplementary Fig. 6a - EZH1/2:62.5%, SIRT:46.2%, JAK: 37.2), indicating similar MIEL profiles across all tested drug concentrations. However, Pan HDAC and HDAC Class I inhibitors displayed progressive profile changes, allowing DA to separate the different concentrations at higher accuracy (Fig. 2a – HDAC Pan: 80.9%; Supplementary Fig. 6a - HDAC Class I: 82.2%).

**Fig. 2:**
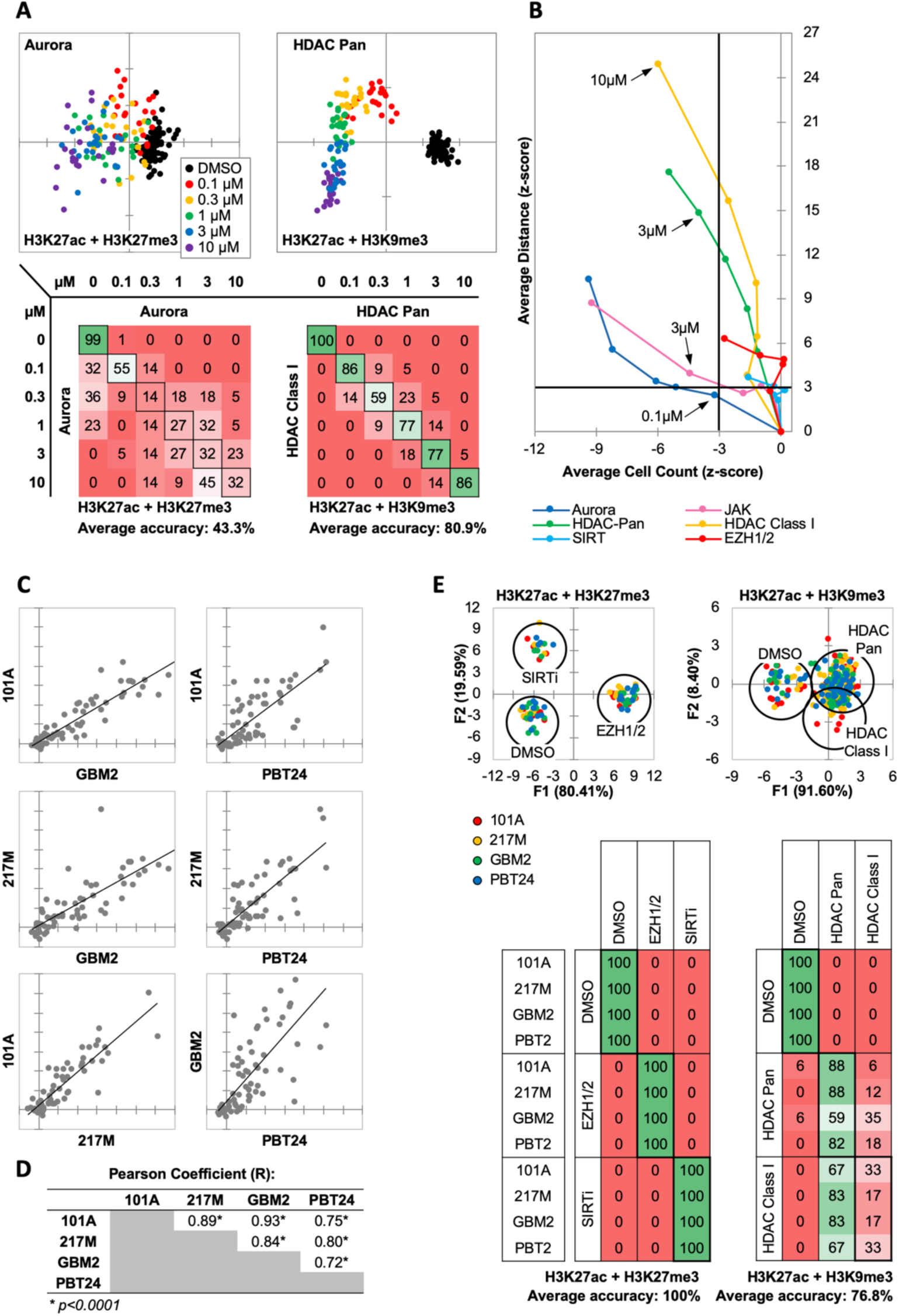
MIEL distinguishes between multiple categories of epigenetic drugs. (A) Quadratic discriminant analysis using texture features derived from images of GBM2 cells treated with DMSO, 0.1, 0.3, 1, 3 or 10 uM Aurora kinase (n=11 compounds, 2 replicates) or HDAC Pan inhibitors (n=11 compounds, 2 replicates) stained for either H3K9me3+H3k27ac or H3K27me3 + H3K27ac. Scatter plots depict the first 2 discriminant factors for each cell population (drug replicate). Confusion matrixes showing results for the discriminant analysis. Numbers represent the percent of replicates classified correctly (diagonal) and incorrectly (off the diagonal). (B) Scatter plot comparing the magnitude of effect (average z-scored Euclidean distances from DMSO) to drug-induced cytotoxicity (average z-scored cell count). Euclidean distance was calculated using image texture features derived from images of H3K27ac + H3K27me3 (Aurora, JAK, SIRT, EZH1/2) or H3K27ac + H3K9me3 (HDAC Pan, HDAC Class I). Distances and cell counts represent average of all compounds in each category; n_Aurora_=11, n_EZH1/2_=5, n_HDAC Class I_=7, n_HDAC Pan_=43, n_JAK_=15, n_SIRTi_=4). (C) Scatter plots comparing the z-scored Euclidean distances from DMSO replicates across 4 GBM lines (n=57 compounds, z-score for each compound is the average of 3 technical replicates). Euclidean distances were calculated using image texture features derived from images of H3K27ac & H3K27me3 or H3K27ac & H3K9me3. (D) A table summarizing the Pearson coefficient and statistical significance of z-scored Euclidean distances shown in “C.” (E) Quadratic discriminant analysis using texture features derived from images of GBM2, PBT24, 101A, 217M cells treated with either DMSO, 5 EZH1/2 inhibitors, 3 SIRT inhibitors, 6 Class I HDAC inhibitors or 17 Pan HDAC inhibitors. Features derived from images of cells stained for H3K27me3 + H3K27ac (EZH1/2, SIRT) or H3K27ac + H3K9me3 (HDACi). Scatter plots depicting the first 2 discriminant factors for each cell population (2 replicates per drug per cell line) color coded according to cell line. Confusion matrix showing classification results for the discriminant analysis (test set, 1 replicate per drug per cell line). Numbers represent the percent of compounds classified correctly (diagonal) and incorrectly (off the diagonal).

In addition to their on-target effect, the drugs may induce epigenetic alterations through toxicity and stress. To estimate the impact of toxicity on changes to drug-induced profiles and its contribution to drug misclassification across a range of concentrations, we plotted z-scored distance from DMSO (effect size) against z-scored nuclei count (a proxy for cytotoxicity) for GBM2 cells treated at a range of drug concentrations (0.1, 0.3, 1, 3, 10µM). This demonstrated that some compound classes, such as Aurora and JAK inhibitors, induce epigenetic alterations only in concentrations where cell count is significantly reduced, whether through toxicity or direct effect on proliferation (Fig. 2b – dark blue and pink respectively), while other compounds, such as HDAC inhibitors, characteristically have a concentration range where epigenetic alterations are not accompanied by reduced cell counts (Fig. 2b – green and yellow). Interestingly, both SIRT and EZH1/2 (Fig. 2b – light-blue and red, respectively) inhibitors affected significant epigenetic changes without inducing significant changes in cell count.

These results indicated the MIEL platform is ideally positioned to analyze dose-dependent effects from drug treatment. In particular, our data suggest that low (0.1uM) and high (10uM) concentration of HDAC inhibitors resulted in distinct and separable epigenetic landscapes, suggesting potentially distinct chromatin/gene expression profiles and divergent biological outcomes when using a low vs high concentration of such compounds.

### MIEL profiles are coherent across multiple cell lines

Testing candidate drugs in multiple cell lines can help gauge their inclusivity and identify tumor subtypes that do not respond to a specific drug or drug class. To test whether MIEL readouts were coherent across multiple glioblastoma TPCs, we treated 4 cell lines with a subset of drugs from the epigenetic library (57 drugs), derived phenotypic profiles, and calculated their effect size (z-scored Euclidean distance from DMSO replicates. This revealed a significant positive correlation between all 4 cell lines pointing to the similarities in their drug sensitivity profiles and demonstrating the robustness of the MIEL read out (Fig. 2c,d). To assess the ability of MIEL to group compounds by function across multiple cell lines we employed DA to classify DMSO and drug treated TPCs across these 4 GBM lines. In this way, we could accurately separate cells treated with drugs modulating distinct functions, such as EZH1/2 or SIRT inhibitors (5 and 3 compounds respectively; mean accuracy 100%; Fig. 2e). However, we were unable to separate drug subclasses with similar functions, such as class I and pan HDACs inhibitors (6 and 17 compounds respectively; mean accuracy 76.8%; Fig. 2e). These results demonstrate MIEL’s ability to correctly categorize by function drugs with varying degrees of potency across multiple cells lines.

Finally, although individual drug activity correlated well across cells lines, the magnitude of the effect for some classes of drugs was highly correlated to the gene expression levels of the target. For example, SIRT inhibition was significantly more effective in lines showing reduced Sirt1 expression (the main SIRT to deacetylate histone 3; n=4 compounds, p<0.02; Supplementary Fig. 6b, c), and there was a significant inverse correlation between Sirt1 expression and the effect size (R=-0.87; Supplementary Fig. 6c).

In sum, we documented that the MIEL assay was both sensitive and robust across multiple primary human glioblastoma cells lines, which further underscores its ability to detect the differences in gene expression and to provide a cumulative measurement of the effect of each compound on cellular epigenetic landscape.

### MIEL helps uncover the mechanism of BET inhibitors synergy with TMZ and ranks their activity

MIEL analysis demonstrated that the magnitude of drug-induced profile change, as measured by the distance from DMSO controls, vary between individual drugs within each drug class (Supplementary Fig. 7a). To test whether these differences are biologically meaningful, we correlated MIEL-based activity of epigenetic drugs which are often designed to work in combination with other treatments (16, 17). One common approach is to use epigenetic drugs to sensitize tumor cells to a standard of care in cytotoxic treatment (15, 43-45), such as radiation and temozolomide (TMZ), which are used to treat glioblastoma. To identify drug classes that sensitize glioblastoma TPCs to cytotoxic therapy, GBM2 cells were treated with epigenetic drugs for 2 days prior to radiation or TMZ. Cytotoxic treatment was carried out for 4 days at levels that induced a 50% reduction in cell numbers (1Gy radiation or 200uM TMZ; Fig. 3a). At the end of the treatment (day 6), cells were counted, and a combined drug index (CDI) was calculated (see Methods). Though we did not identify any drugs that synergized (CDI<0.7) with the radiation therapy (Fig. 3b, right panel), multiple drugs from both PARP and BET inhibitor (PARPi and BETi) sensitized cells to TMZ (Fig. 3b, left panel).

**Fig. 3:**
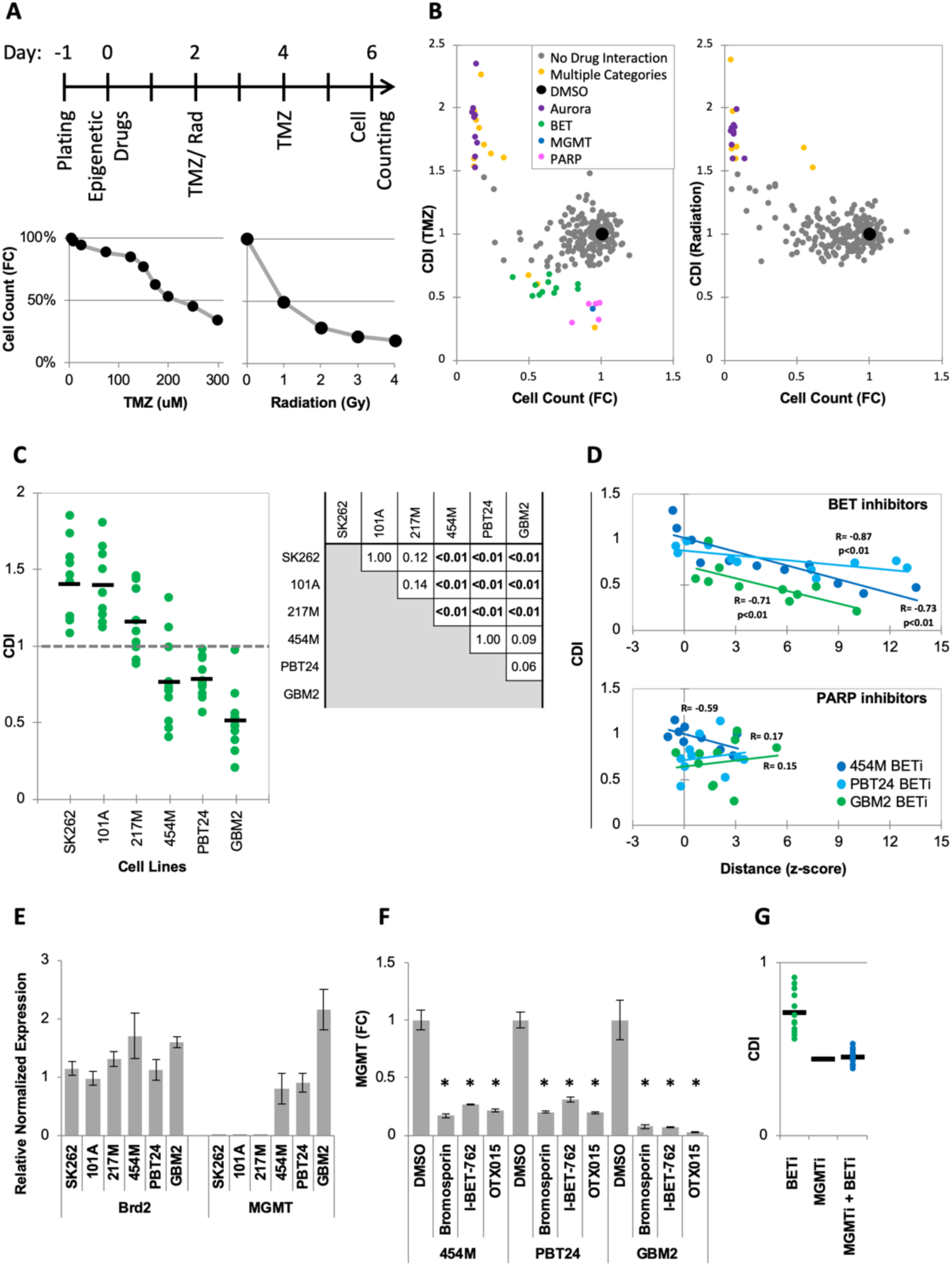
MIEL can be used to rank candidate drugs by activity. (A) Top: Scheme describing the experimental setup used to identify synergy between epigenetic drugs and radiation or TMZ. Bottom: Scatter plots showing the fold reduction in GBM2 cell count following a 4-day treatment with varying TMZ concentration and radiation doses. (B) Scatter plots showing fold change in cell count (compared to DMSO treated cells) and coefficient of drug interaction (CDI) for synergy with TMZ (left) and radiation (right) for each drug (n=222, values represent the average of 3 technical replicates). (C) Graph showing individual and average CDI for BET inhibitors in 6 GBM lines (n=11 drugs, average of 3 technical replicates; p-values calculated by ANOVA using Tukey’s HSD for multiple comparisons between lines and shown in table). (D) Scatter plot showing the correlation between CDI and MIEL-derived activity (z-scored Euclidean distance from DMSO) of BET and PARP inhibitors (n_BETi_=11; n_PARPi_=10; values represent the average of 3 technical replicates) in 3 GBM lines (454M, PBT24, GBM2). (E) Bar graph showing the relative normalized expression of Brd2 and MGMT in 6 GBM lines as measured by qPCR (Mean±SD; n=3 technical repeats). (F) Bar graph showing fold reduction in MGMT expression following treatment with BET inhibitors in 3 different GBM lines as measured by qPCR (Mean±SD; n=3 technical repeats). (G) Graph showing individual and average TMZ sensitization CDI for BETi, MGMTi (Lomeguatrib) and BETi & MGMTi in GBM2 cells (n=11 drugs, values represent the average of 3 technical replicates).

PARPi have been extensively studied in this context and have been shown to function through multiple nonepigenetic mechanisms such as PARP trapping (46–48). Consistent with this, most PARPi did not induce detectable epigenetic changes using MIEL (Fig. 3d, Supplementary Fig. 7b), and we found no correlation between the magnitude of epigenetic changes as measured by MIEL and CDI (Fig. 3d – bottom panel). To date, only a single report utilizing the BETi OTX015 (49) has pointed to synergy with TMZ, prompting us to validate this finding in 6 additional glioblastoma lines. In 3 lines, BETi increased the TMZ effectiveness (average CDI: 454M 0.76±0.28, PBT24 0.78±0.12 and GBM2 0.51±0.2; Mean±SD; n=11 BETi; Fig. 3c). In the other 3 lines, the drugs did not synergize and, in many cases, were found to be protective against (CDI>1) TMZ (average CDI: SK262 1.4±0.26, 101A 1.4±0.22 and 217M 1.2±0.21; Mean±SD; n=11 BETi; Fig. 3c; p-values for all pairwise comparisons). Only a few BETi-induced epigenetic changes occurred during our 24-hour initial screening (Fig. 1b). However, following 6 days of treatment, 6 of 11 BETi induced significant (average z-scored distance from DMSO replicates >3) epigenetic changes in all cell lines (Fig. 3d, Supplementary Fig. 7b). In lines displaying TMZ and BETi synergy, the degree of BETi activity, as measured by MIEL, significantly correlated with the degree of synergism (Fig. 3d – top panel). This demonstrated that for individual compounds, MIEL can predict relative drug activity and suggests an epigenetic component for the mechanism of BETi-TMZ synergy.

O^6^-alkylguanine DNA alkyltransferase (MGMT), which provides the main line of defense against DNA alkylating agents such as TMZ, has been found to be epigenetically silenced through DNA methylation in a large fraction of glioblastoma tumors (50, 51). To gain a better understanding of the mechanism by which BETi sensitizes glioblastoma TPCs to TMZ treatment, we quantified MGMT expression in the 6 lines tested using qPCR. Analysis showed that while all lines expressed similar BET-TF levels, such as Brd2 (Fig. 3e), and were thus susceptible to BET inhibitors, only 3 lines displaying BETi-TMZ synergy expressed MGMT (Fig.3e). Yet after treating those 3 lines with BETi, MGMT expression was dramatically reduced (Fig. 3f). Finally, after combining BET inhibitors with the MGMT inhibitor Lomeguatrib, we detected no increase in sensitivity to TMZ above the levels conferred by Lomeguatrib alone (Fig. 3g).

In sum, we have discovered that several BETi synergized with TMZ treatment by reducing MGMT expression. We determined that the degree of synergism displayed by individual BETi positively correlated with the magnitude of their epigenetic effect as measured using MIEL, suggesting that their mechanism of action involves epigenetic change. In contrast, the activity of PARP inhibitors didn’t correlate with MIEL distance, suggesting an alternative mechanism of action unrelated to epigenetic changes.

### MIEL discriminates between multiple cell fates

To determine whether MIEL could discriminate between different cell fates we analyzed 3 cell types: primary human fibroblasts, induced pluripotent stem cells (iPSCs) derived from these fibroblasts, and neural progenitor cells (NPCs) differentiated from the iPSCs. The fibroblasts were isolated from 3 unrelated donors (WT-61, WT-101, WT-126) and used to obtain corresponding iPSC and NPC lines. Cellular identities of the 3 cell types were verified by immune-fluorescence for Sox2 and Oct4 (Fig. 4a), and MIEL analysis was carried out using data from either H3K4me1 and H3K9me3 or H3K27ac and H3K27me3 staining, with both combinations providing similar results. Multivariate centroids were calculated for each cell population and plotted on a distance map to visualize the relative Euclidean distance between various cell populations. The fibroblasts, iPSCs, and NPCs each segregate to form 3 visually distinct territories (Supplementary Fig. 8a, c). We separated the 9 lines by cell-fates using DA, which showed an accurate separation of the different cell-fates across all 3 donors (average accuracy 100%; Fig. 4b, Supplementary Fig. 8e). A similar analysis performed to separate the different donors showed only low accuracy (average accuracy 55.5%; Fig. 4c, Supplementary Fig. 8f). To determine whether it was possible to discriminate between individual cells with different fates, a Support Vector Machine (SVM) classifier was trained on a subset of fibroblasts, iPSCs, and NPCS from the 3 donors. Classification of the test set indicated a high degree of separation between the different fates at a single cell level (Supplementary Fig. 8b, d). Additionally, MIEL analysis (using only H3K9me3) was able to discriminate between the main lineages of primary hematopoietic cell types freshly isolated from mouse bone marrow, namely lymphoid, myeloid, and stem/progenitors (Supplementary Fig. 9). However, closely related cell types in each lineage such as hematopoietic stem and progenitor cells were not readily separated (Supplementary Fig. 9).

**Figure 4.**
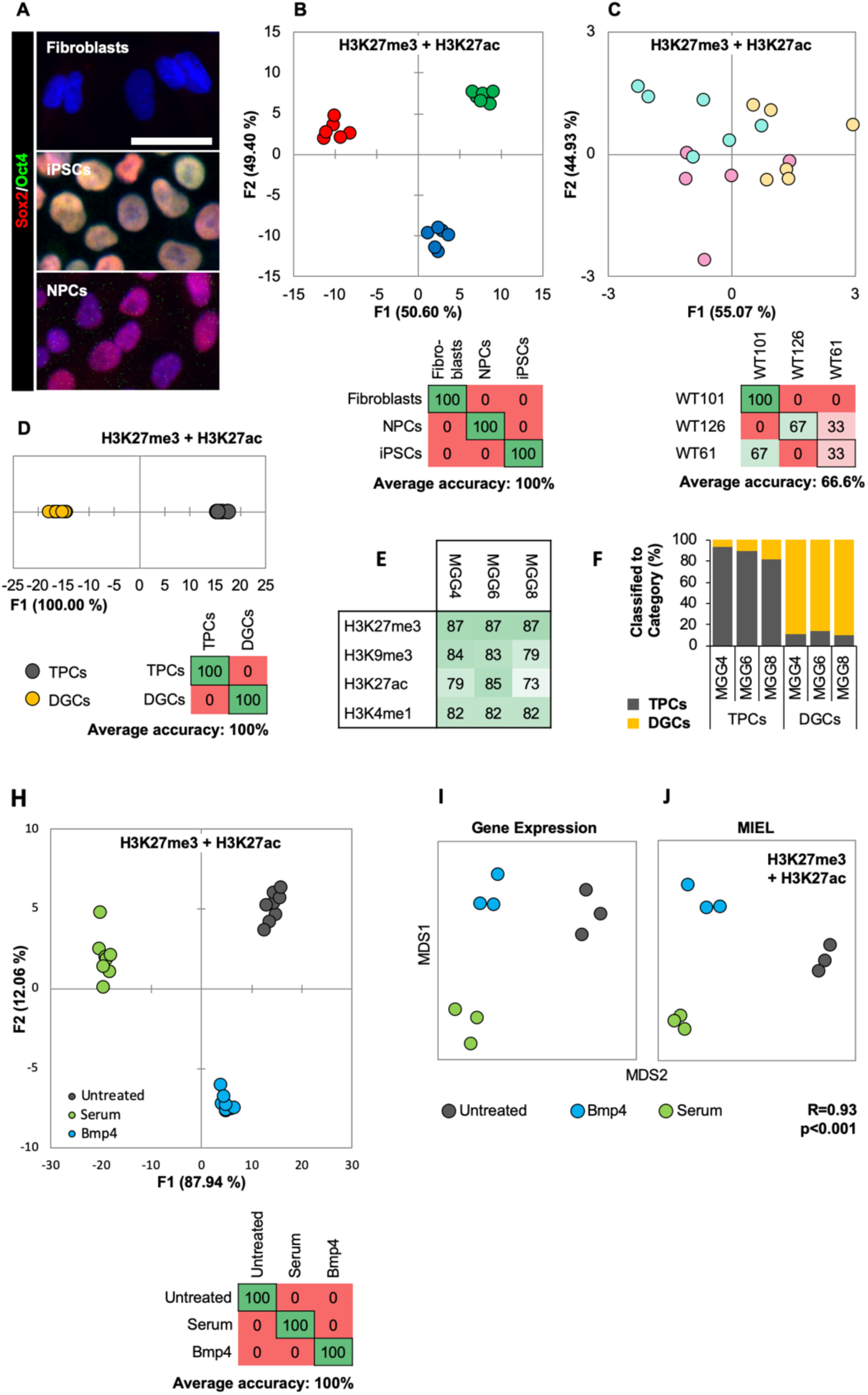
MIEL can distinguish between cell fates and identify glioblastoma differentiation. (A) Hoechst 33342 stained (blue), and Sox2 (red) and Oct4 (green) immunofluorescence labeled fibroblasts (Sox2^-^/Oct4^-^), iPSCs (Sox2^+^/Oct4^+^) and NPCs (Sox2^+^/Oct4^-^). Scale bar, 50 µm. (B, C) Quadratic discriminant analysis separating either cell fates or cell lines using texture features derived from images of fibroblasts, iPSCs, and NPC lines from 3 human donors (WT-61, WT-101 and WT-126; 3 technical replicates each); stained for H3K9me3 and H3K4me1. (B) Discriminant analysis separating the different cell types. Scatter plot depicts the first 2 discriminant factors for each cell population (2 replicate per cell line and cell type). Confusion matrix showing classification results for discriminant analysis (test set: 1 replicate per cell line and cell type). Numbers represent the percent of correctly (diagonal) and incorrectly (off the diagonal) classified cell populations. (C) Discriminant analysis attempts to separate different cell lines. Scatter plot depicts the first 2 discriminant factors for each cell population (2 replicates per cell line and cell type). Confusion matrix showing classification results for discriminant analysis (test set: 1 replicate per cell line and cell type). Numbers represent the percent of correctly (diagonal) and incorrectly (off the diagonal) classified cell populations. (D, E, F) TPC and DGC cell lines derived simultaneously from tumors of 3 human donors (MGG4, MGG6, MGG8; 3 technical replicates each); stained for H3K9me3, H3K4me1. (D) Quadratic discriminant analysis separating TPCs and DGCs using image texture features. Scatter plot depicts the first discriminant factor for each cell population (2 replicates per cell line). Confusion matrix showing results of discriminant analysis (test set: 1 replicate per cell line). Numbers represent the percent of correctly (diagonal) and incorrectly (off the diagonal) classified cell populations. (E) Pairwise classification of single TPC and DGC cells using an SVM classifier trained on texture features derived from images of H3K27me3, H3K9me3, H3K27ac, or H3K4me1. Numbers correspond to the percent of correctly classified cells for each line using indicated epigenetic marks. (F) Bar graph shows results of SVM classification for single TPC and DGC cells using a classifier trained on texture features derived from images of H3K27ac and H3K27me3 marks in the MGG4 line. (H) Quadratic discriminant analysis using texture features derived from images of untreated or 2 days serum or Bmp4 treated GBM2, 101A, SK262 and 454M cells (3 replicates per cell lines per treatment) and stained for H3K9me3, H3K4me1. Scatter plot depicts the first 2 discriminant factors for each cell population (2 replicates per cell lines per treatment). Confusion matrix shows classification results for discriminant analysis (test set: 1 replicate per cell line per treatment). Numbers represent the percent of correctly (diagonal) and incorrectly (off the diagonal) classified cell populations. (I) Distance map depicting the relative Euclidean distance between the transcriptomic profiles of DMSO-, Bmp4- and serum-treated GBM2 cells calculated using FPKM values of all expressed genes (14,376 genes; FPKM>1 in at least one sample). Each treatment in triplicates. (J) Distance map depicting the relative Euclidean distance between the multiparametric centroids of DMSO-, Bmp4- and serum-treated GBM2 cells calculated using texture features derived from images of H3K27ac and H3K27me3 marks. Each treatment in triplicates. R denotes Pearson correlation coefficient.

These results underscore MIEL’s ability to discriminate multiple different cell types and differentiation states uniquely based on their single-cell epigenetic landscapes both in cultured and primary cells of human and mouse origin.

### MIEL determines the signatures of glioblastoma stem cells and differentiated glioblastoma

Most epigenetic drugs are known to directly affect the level histone and DNA modifications, which are the substrates MIEL assay. To test whether MIEL is capable to identify and classify drugs that affect epigenetic landscape indirectly, we focused on glioblastoma differentiation paradigm. Although such approach was proposed by several groups (24–26), identification of small molecule inducers of glioblastoma differentiation has been challenging. Previous attempts to design screening strategies for this purpose have met with multiple difficulties. One critical problem is the lack of informative markers faithfully reporting GBM differentiation that could be used for high-throughput screening (10). Therefore, we tested the utility of MIEL platform to screen for drugs inducing glioblastoma TPCs differentiation.

We tested MIEL’s ability to distinguish TPCs and differentiated glioma cells (DGCs), derived from primary human glioblastomas (10). Three TPC/DGC pairs were derived in parallel from 3 genetically distinct glioblastoma tumor samples (MGG4, MGG6, and MGG8) over a 3-month period using either serum-free FGF/EGF for TPCs or 10% serum for DGCs (10). Visualization using a distance map demonstrated that TPCs and DGCs segregate to form two visually distinct territories (Supplementary Fig. 8g) and were separated with high accuracy using DA (mean accuracy 100%; Fig. 4d). SVM-based pairwise classification of single cells distinguished TPCs from their corresponding DGC lines with an average accuracy of 83%, using any of the 4 epigenetic marks tested (H3K27me3, H3K9me3, H3K27ac, and H3K4me1; Fig. 4e). An SVM classifier derived from images of the MGG4 TPC/DGC pair separated all 3 TPC/DGC pairs with 88% average accuracy, providing proof of principle for the derivation of a signature for nontumorigenic cells obtained following serum differentiation of primary glioblastoma cells (Fig. 4f).

These findings suggest that MIEL can readily distinguish undifferentiated TPCs from differentiated DGCs based on multiparametric signatures of these glioblastoma cells using only the patterns of universal epigenetic marks. Of note, until now such signatures could only be obtained using simultaneous assessment of dozens of transcripts by averaging thousands of cells (10, 52).

### Short-term treatment with serum or Bmp4 initiates TPC differentiation

For the purpose of establishing a screening protocol, we tested whether short serum or Bmp4 treatment is sufficient to induce a differentiation-like phenotype in TPCs. We treated several glioblastoma cell lines for 3 days with either serum or Bmp4, then quantified expression of core transcription factors previously shown to determine the TPC transcriptomic program (10). Immunostaining revealed that the 4 transcription factors Sox2, Sall2, Brn2 and Olig2 were downregulated by both serum and Bmp4 in a cell line-dependent manner (Supplementary Fig. 10a). RNAseq analysis of serum- and Bmp4-treated GBM2 cells revealed that 3 days of treatment reduced (vs untreated cells) expression of most genes previously found to constitute the transcriptomic stemness signature (52) (Supplementary Fig. 10b). Additionally, both serum and Bmp4 were found to attenuate TCP growth rate (Supplementary Fig. 10c). To identify the cellular processes altered by these treatments, we conducted differential expression analysis. Expression of 4852 genes was significantly altered (p<0.01 and −1.5<FOLD Change >1.5) by either serum or Bmp4. Gene Ontology (GO) analysis of these altered genes indicated enrichment in multiple GO categories consistent with initiation of TPC differentiation; these include cell cycle, cellular morphogenesis associated with differentiation, differentiation in neuronal lineages, histone modification, and chromatin organization (Supplementary Fig. 11).

These results demonstrate that a 3-day treatment with either serum or Bmp4 is sufficient to induce transcriptomic changes characteristic of TPC differentiation. Previous work indicated distinct features of glioblastoma differentiation induced with BMP compared to serum (27). Indeed, we observed distinct expression changes, including differences in expression of genes regulating chromatin organization and histone modifications (Supplementary Fig. 12a, b), between serum- and Bmp4-induced glioblastoma differentiation.

### MIEL detects epigenetic changes following short-term serum or Bmp4 treatment

We treated 4 genetically distinct glioblastoma lines with serum or BMP4, then conducted MIEL analysis using either H3K9me3 and H3K4me1 or H3K27ac and H3K27me3 to detect TPC differentiation. Discriminant analysis allowed high accuracy separation of these treatments across all cell lines using both histone modification combinations (mean accuracy 100%; Fig. 4h; Supplementary Fig. 12c).

The global gene expression profile represents a gold standard for defining the cellular state (53). To test whether MIEL reliably reports the epigenetic changes associated with serum and Bmp4 treatments we conducted a correlation between MIEL-based and global gene expression-based metrics. We sequenced untreated and 3 days serum- or Bmp4-treated GBM2 TPCs. All genes with FPKM>1 in at least one cell population were used to calculate the Euclidean distance matrix between all cell populations. FPKM-based distances were then correlated to image texture feature-based distances. The resulting Pearson correlation coefficient of R=0.93 (p<0.001) suggests a high correlation between these two metrics (Fig. 4j, k), demonstrating that MIEL is capable of distinguishing closely related glioblastoma differentiation routes induced by serum or BMP and validating the robustness of the MIEL approach for analyzing glioblastoma differentiation.

### MIEL successfully prioritizes small molecules inducing TPCs differentiation

We screened the Prestwick compound library (~1200 compounds) using MIEL to identify compounds inducing glioblastoma TPC differentiation based on the differentiation signatures obtained with serum/Bmp4 treatments. GBM2 TPCs were treated for 3 days with Prestwick compounds at 3 µM fixed, then immunolabeled for H3K27ac and H3K27me3. GBM2 cells treated with DMSO, serum, BMP4, or compound were compared within the same plate (to avoid imaging artifacts and normalization issues).

To identify epigenetically active compounds, we calculated the Euclidean distance to the DMSO center for each DMSO replicate and Prestwick compound. Distances were z-scored, and active compounds were defined as compounds for which z-scored distance was greater than 3. Compounds with less than 50 cells imaged were considered toxic and excluded from analysis. Following analysis, MIEL identified 144 active compounds. To identify compounds inducing epigenetic changes reminiscent of serum-BMP4-induced differentiation, we used quadratic DA to build a model separating untreated, serum-treated, and Bmp4-treated cells and classified all 144 active compounds to these categories (Fig. 5a,b). A total 31 compounds were classified as similar to either serum or Bmp4 (i.e., differentiated). Of these, 20 compounds belonged to 1 of the following 4 categories: Na/K-ATPase inhibitors of the digoxin family, molecules that disrupt microtubule formation or stability, topoisomerase inhibitors, or nucleotide analogues that disrupt DNA synthesis (Fig. 5b). To further narrow down the list of candidates, we conducted pairwise SVM classification of DMSO- and either serum- or BMP4-treated cells, then selected compounds that induced at least 50% of the cells to be classified as either serum- or BMP4-treated. We then calculated the Euclidean distance between candidate compounds and serum- and BMP4-treated cells; we selected compounds where the distance to one or both treatments was less than the distance between DMSO and that treatment. Of the 20 candidate compounds identified, 15 belonged to 1 of the 4 categories mentioned above (Supplementary Fig. 13a). From the 15 candidate compounds, we chose 2 top compounds from each of the four categories (8 total) for further analysis. GBM2 cells were treated for 3 days with DMSO, serum, Bmp4 or candidate compounds at 0.3,1, or 3 µM, fixed, then immunostained for H3K27ac and H3K27me3. Using pairwise SVM-based classifications of untreated cells and either serum- or Bmp4-treated cells identified for each of the 8 compounds, the lowest concentration at which at least 50% of the cells were classified as treated (Supplementary Fig. 13b) and used those concentrations for all subsequent experiments. Because most of these compounds are known for their cytotoxic effects, we verified the growth rates of drug-treated glioblastoma cells. With the exception of digoxin, which was cytostatic, drug treatment resulted in growth rates comparable with those induced by serum or BMP4 (supplementary Fig. 14a). We used immunofluorescence to test for expression of core TPC transcription factors (Sox2, Sall2, Brn2 and Olig2). Except for trifluridine, all compounds induced statistically significant reductions in Sox2; digoxin and digitoxigenin also induced a significant reduction of Sall2 and Brn2; olig2 expression was unaltered by any treatment (Supplementary Fig. 14b).

**Figure 5.**
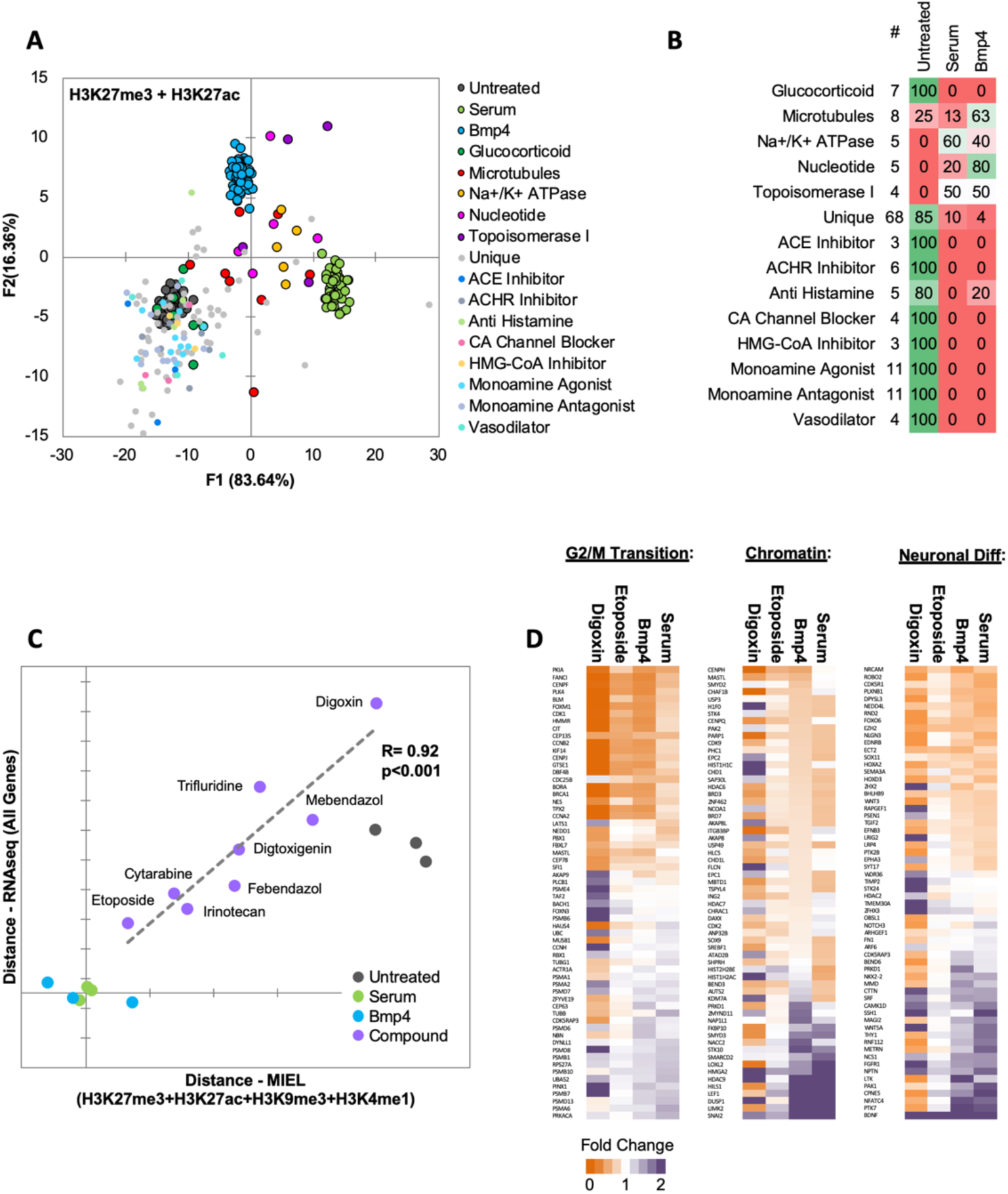
MIEL prioritizes small molecules based on serum/Bmp4 differentiation signature. (A) Scatter plot depicting the first 2 discriminant factors for untreated and serum-, Bmp4- and compound-treated cells (1 replicate per compound). Quadratic discriminant analysis using texture features derived from images of untreated and serum-, Bmp4- and compound-treated GBM2 cells stained for H3K27me3, H3K27ac. Model was built to separate untreated and serum- and Bmp4-treated cells (60 technical replicates each). (B) Confusion matrix shows classification of epigenetically active Prestwick compounds. Numbers depict the percent of compounds from each category classified as either untreated, serum or Bmp4 treated. (C) Scatter plot shows the correlation of gene expression profile-based ranking and MIEL-based ranking for 8 candidate drugs, untreated, serum- or Bmp4-treated GBM2 cells. Euclidean distance to serum- or Bmp4-treated GBM2 cells was calculated using transcriptomic profiles (vertical axis) or texture features derived from images of H3K27ac and H3K27me3, H3K9me3, and H3K4me1 marks (horizontal axis). Distances were normalized to untreated and serum- or Bmp4-treated GBM2 cells. (D) Heat maps showing fold change in expression of select genes taken from the Gene Ontology (GO) list: cell cycle G2/M phase transition (GO:0044839), chromatin modification (GO:0006325), and regulation of neuron differentiation (GO:0045664). R denotes Pearson correlation coefficient. Drug concentrations a-c: febendazole=0.5 µM, mebendazole=0.5 µM, cytarabine=0.3 µM, trifluridine=3 µM, irinotecan=0.5 µM, etoposide=0.3 µM, digitoxigenin=0.3 µM, digoxin=0.3 µM.

Next, we investigated whether the compounds identified using MIEL can induce transcriptomic changes similar to serum and Bmp4 treatment and quantified the ability of MIEL to predict compounds best at mimicking these treatments. GBM2 cells were treated with DMSO, serum, Bmp4, or each of the eight candidate compounds; after 3 days, RNA was extracted and sequenced. Transcriptomic profiles of the eight compounds were ranked according to average Euclidean distance (based on FPKM values for all expressed genes) from serum- or BMP4-treated cells. To safeguard against potential artefacts of cytotoxicity, we compared gene expression-based ranking with measured cellular growth rates from drug treatments and found no positive correlation (Supplementary Fig. 14c). When we next compared Sox2 expression levels under all treatment conditions to determine whether the transcription factor can identify drugs that best mimic serum or BMP4, we found no positive correlation between either expression levels or transcriptomic-based rankings (Supplementary Fig. 14d), suggesting that Sox2 levels alone are insufficient to stratify the compounds. Finally, to compare MIEL-based signatures to the transcriptomic profile, we ranked MIEL readouts of cells treated with the eight drugs according to average Euclidean distance from serum- or Bmp4-treated cells (calculated using texture features derived from images of H3K27ac, H3K27me3, H3K9me3, and H3K4me1). Comparison of the MIEL-based metric with the gene expression-based metric revealed a high degree of positive correlation between MIEL- and gene expression-based rankings (Pearson correlation coefficient R=0.92, p<0.001; Fig. 5c). To further visualize these results, we constructed heat maps depicting fold change in gene expression associated with several GO terms enriched by serum and Bmp4. Our top candidate, etoposide, altered expression of a large portion of genes in similar fashion to that of serum and BMP4; in contrast, the lowest-ranking candidate, digoxin, induced changes in gene expression, which were rather different from serum and BMP4 (Fig. 5d).

Taken together, the above results suggest unique ability of MIEL to identify molecules that shift epigenetic signature of glioblastoma TPCs towards DGCs. Critically, MIEL is capable of ranking such molecules according to their change-inducing potency and that ranking robustly correlate with global expression-based readouts of glioblastoma differentiation.

## Discussion

Here we have introduced MIEL, a novel method that expands phenotypic profiling to take advantage of universal biomarkers present in all eukaryotic cells, histone modification, and exploits the patterns of chromatin organization and epigenetic marks. The pipeline we developed employs information extracted from immunofluorescence images of specific histone modifications and is geared towards drug discovery and high-throughput screening. Focusing on compounds that modulate epigenetic writers, erasers, and readers, we have shown that MIEL markedly improves detection compared to conventional intensity-based thresholding approaches and enables functional categorization of such compounds. We have demonstrated that MIEL readouts are coherent across multiple compound concentrations and cell lines and can provide information regarding drug activity levels and their mechanism of action. We have also documented MIEL ability to robustly report cellular fate and provide proof of concept for identifying and prioritizing drugs inducing differentiation of glioblastoma TPCs.

Previous studies have demonstrated that image-based profiling can distinguish between classes of compounds with very distinct functions, such as Aurora and HADC inhibitors (5). One objective of our study was to estimate the resolution of separation between categories of compounds with similar functions. We found that a single histone modification was sufficient to separate highly distinct classes. Separating similar classes (e.g., Aurora and JAK inhibitors, which affect histone phosphorylation, or Pan and Class I HADCs, which affect histone acetylation) required staining for at least one additional histone modification. Despite their many advantages, cellular assays, including high-content assays, are often used as secondary screens for epigenetic drugs due to multiplicity of enzyme family members and an inability to determine direct enzymatic activity (54). Consequently, MIEL’s ability to separate closely related functional categories on top of other advantages make this profiling approach an attractive alternative for primary screens.

Phenotypic profiling methods have been previously used to identify genotype-specific drug responses by comparing profiles across multiple isogenic lines (55). Here we show that biologic activity (i.e., serum and Bmp4) that induces glioblastoma differentiation, as well as that of 57 epigenetic compounds, was significantly correlated across four different primary glioblastoma lines. We also showed that variation in activity levels correlated with target expression levels and that the various categories can be distinguished across cell lines. Together, these suggest that MIEL could be used to identify cell lines showing an aberrant reaction to selected drugs and, therefore, aid in identifying optimal treatments for individual patients. Similar applications have previously been used to tailor specific kinase inhibitors to patients with chronic lymphocytic leukemia (CLL) who display venetoclax resistance (56).

Given the limited success of cytotoxic drugs to treat glioblastoma, we focused on alternative approaches: (1) epigenetic drugs aimed at sensitizing glioblastoma TPCs to such treatments, and (2) inducing glioblastoma differentiation. We have demonstrated MIEL’s ability to rank candidate drug activity to correctly predict the best candidates for achieving the desired effect. The importance of this is highlighted in large (hundreds of thousands of compounds) chemical library screens, which typically identify many possible hits needing to be reduced and confirmed in secondary screens (57, 58).

Our results uncovered a strong correlation between BET inhibitor activity (measured by MIEL) and its ability to synergize with TMZ and reveal a previously unknown role for BET inhibitors in reducing MGMT expression. Previous studies have demonstrated upregulation of several BET transcription factors in glioblastomas (59, 60), and multiple pre-clinical studies have investigated the potential of BET inhibition as a single drug treatment for glioblastoma (61–63). However, while clinical trials with the BET inhibitor OTX015 demonstrated low toxicity at doses achieving biologically active levels, no detectable clinical benefits were found (64). This prompted approaches using drug combinatorial treatments (65) such as combining HDACi and BETi (66, 67). However, the mechanism by which BETi induces increased TMZ has not been described. Recently, a distal enhancer regulating MGMT expression was identified (68). Activation of this enhancer by targeting a Cas9-p300 fusion to its genomic locus increased MGMT expression while deletion of this enhancer reduced MGMT expression (68). As BET transcription factors bind elevated H3K27ac levels found in enhancers (69, 70), this may be a possible mechanism for BETi-induced reduction of MGMT expression, which in turn result in increased sensitivity to the DNA alkylating agent TMZ.

Silencing the MGMT gene through promoter methylation has long been known to make TMZ treatment more responsive and to improve prognosis in patients with glioblastoma (50, 51, 71). Yet, clinical trials that combine TMZ and MGMT inhibitors have not improved therapeutic outcomes in such patients, possibly due to the 50% reduction in dose of TMZ, which is required to avoid hematologic toxicity (72–74). Thus, BETi offers an attractive line of research, though further studies are needed to determine whether the elevated sensitivity of glioblastoma to BETi, and its ability to reduce MGMT expression, thus synergizing with TMZ, could be exploited to improve patient outcome.

Based on our success with identifying the mechanism of BETi action, we believe that MIEL approach is well positioned to systematically analyze and identify epigenetically active compounds, then provide critical initial information for their mechanism of action.

We previously analyzed serum and BMP4, two established biologicals known to induce glioblastoma differentiation in culture (24–26), and established signatures of the differentiated glioblastoma cells based on the pattern of epigenetic marks that could be applied across several genetic backgrounds. This is the first time that a signature for glioblastoma differentiation suitable for high-throughput drug screening has been obtained. Indeed, results of previous studies using bulk glioblastoma analysis (27) or single-cell sequencing (52) could not be readily applied for high-throughput screening. As a proof of principle, we analyzed the Prestwick chemical library of 1200 approved drugs to validate MIEL’s ability to select and prioritize small molecules, which mimic the effect of serum and BMP4, using global gene expression profiling. Surprisingly, we observed that the degree of reduction in endogenous SOX2 protein levels following drug treatment did not correlate with the degree of differentiation assessed by global gene expression; in contrast, MIEL-based metrics did correlate. This result, taken together with MIEL’s ability to distinguish multiple cells types (iPSCs, NPCs, fibroblasts, hematopoietic lineages) across several genetic backgrounds, suggests that the MIEL approach does not only readily identify compounds by inducing desired changes in cell fate but, specifically, can be a cost-effective tool for prioritizing hundreds of thousands of compounds during the primary screenings.

By tapping into the wealth of information contained within the cellular epigenetic landscape through modern high-content profiling and machine-learning techniques, the MIEL approach represents a valuable tool for high-throughput analytical and drug discovery and is especially relevant when the desired cellular outcome cannot be readily defined using conventional approaches.

## Acknowledgments

We are thankful to Harley Kornblum (UCLA) for sharing multiple primary human glioblastoma lines, Alysson Muotri (UCSD) for providing fibroblast, iPSC and NPC lines, and Bradley Bernstein (MGH Harvard) for sharing MGG-TPCs and MGG-DGCs lines, Laure Escoubet (Celgene) for discussions and support, Alex Kiselyov (Genea Biocells) for help with initial compound libraries and discussions. We owe a debt of gratitude to Susanne Heynen-Genel, Debbie Chen, and other members of the High-Content Facility at CPCCG for their invaluable help with cell imaging and to Brian James and Kang Liu at the SBP Genomics core for their help with library preparation and RNA sequencing (NCI Cancer Center Support Grant P30 CA030199). We thank Linda Penn for suggestions and help with the manuscript. This work was supported by sponsored research agreement with Celgene to A.V.T., an R01 NS066278 to A.V.T, and by a CIHR Foundation grant and a Tier 1 Canada Research Chair award to D.W.A.

---

**Supplementary Fig. 1.**
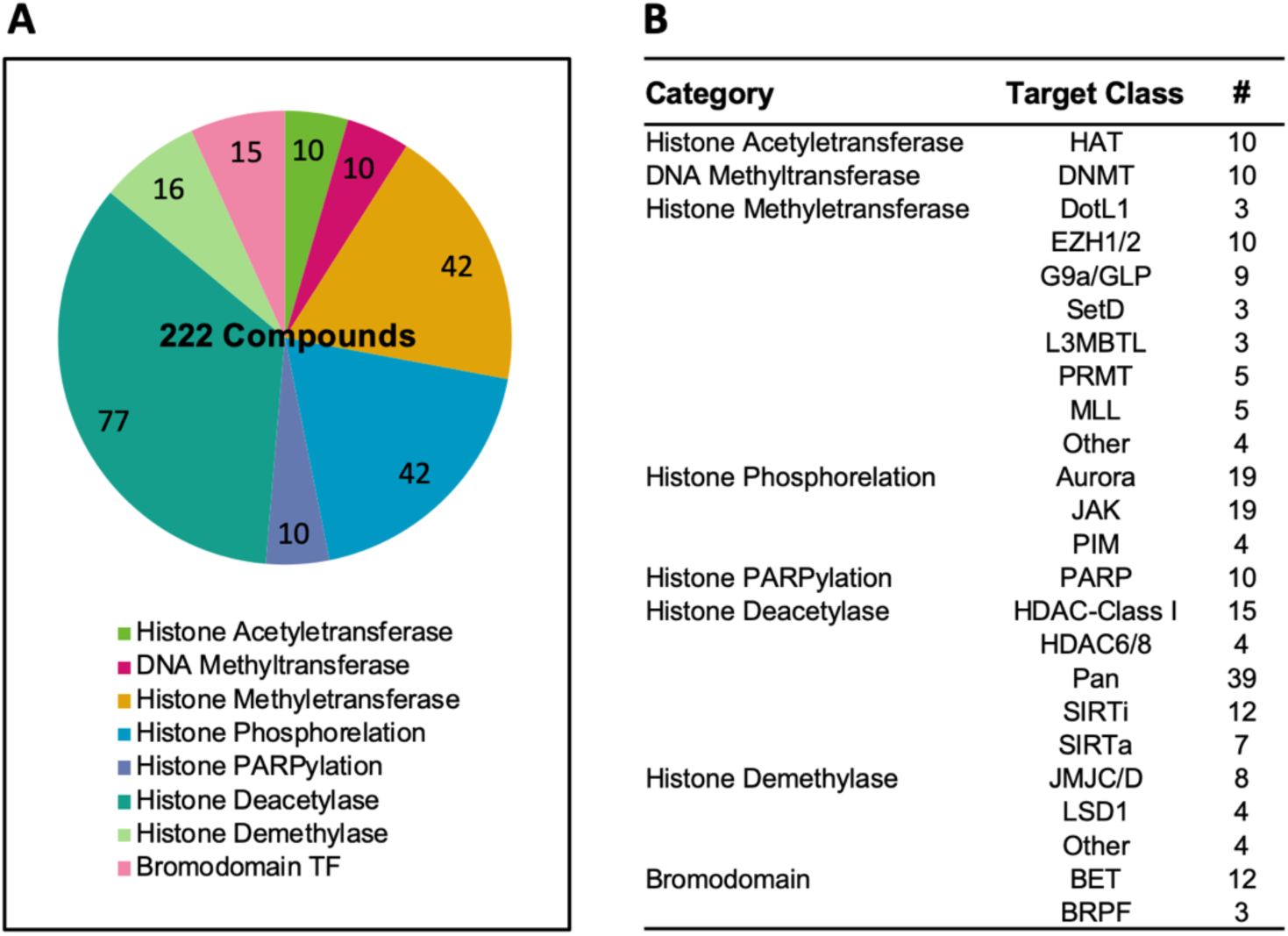
(A) Pi chart showing functional classes of epigenetic drugs used in the study. (B) Table detailing the molecular targets of epigenetic drugs used in the study.

**Supplementary Fig. 2.**
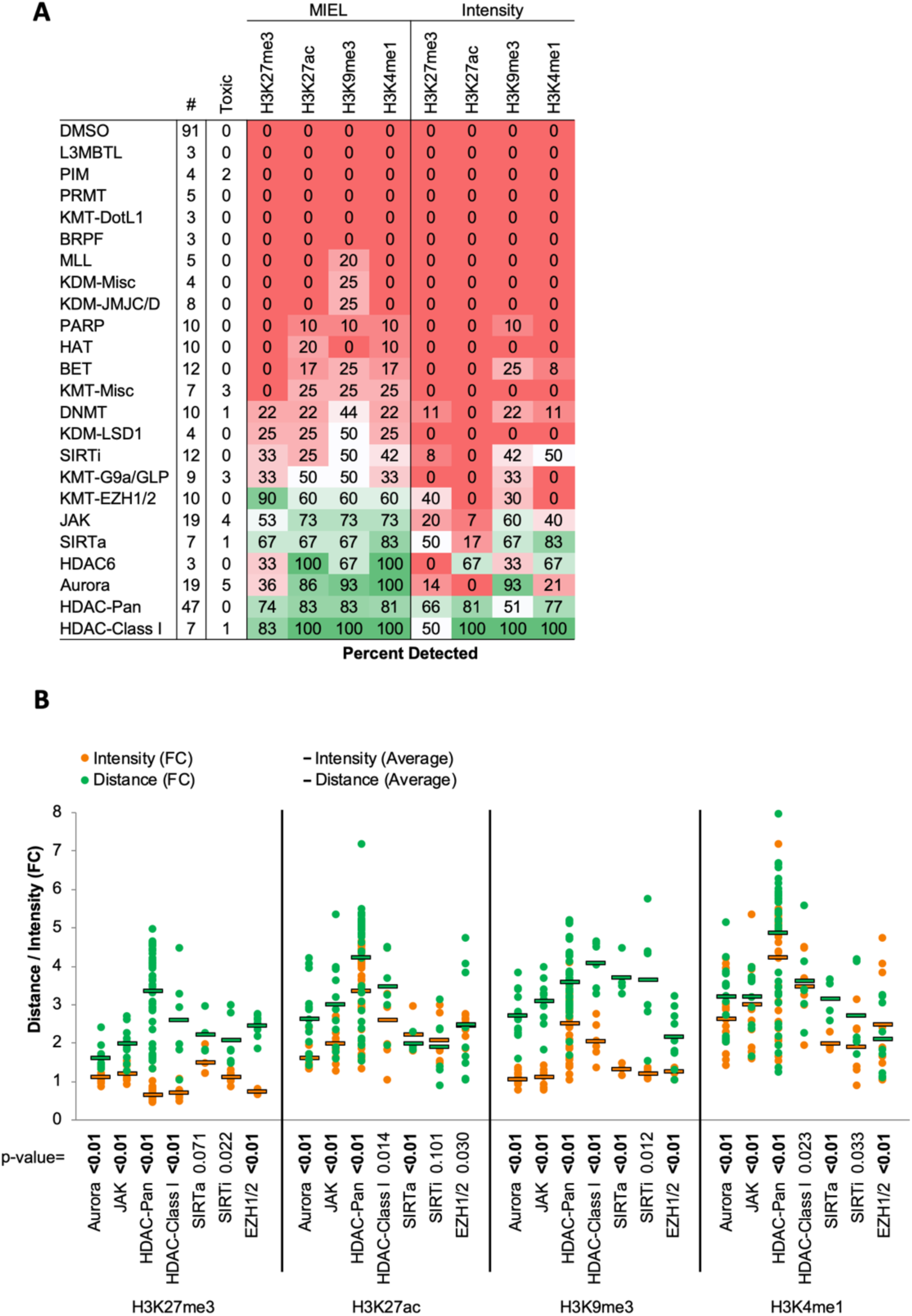
(A) Table showing the fraction of epigenetic drugs in each functional category identified as active by either MIEL analysis employing texture features derived from images of GBM2 cells stained for either H3K9me3, H3K4me1, H3K27ac, H3K27me3 or by intensity-based analysis using individual modifications (see Methods). (B) Bar graph depicting the average fold change in Euclidean distance from DMSO replicates induced by drugs from several functional categories as calculated using mean intensity or using texture features derived from images of individual histone modification (Mean±SD; p-values calculated by ANOVA using Tukey’s HSD for multiple comparisons and shown in table; n for each category is shown in “A”).

**Supplementary Fig. 3.**
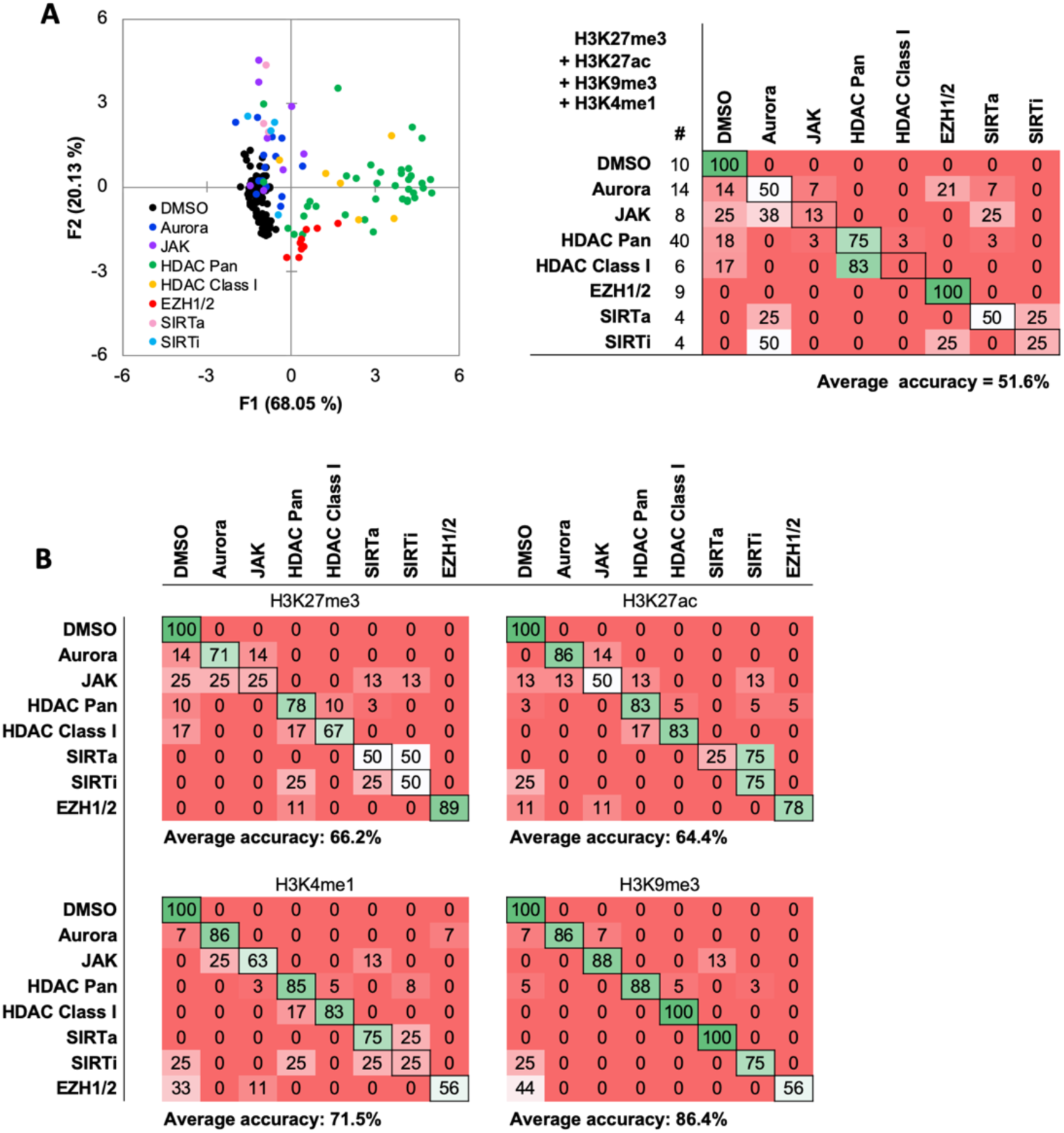
(A) Quadratic discriminant analysis using average fluorescent intensity derived from images of GBM2 cells treated with either DMSO or 85 active compounds (2 technical replicates per compound; 38 DMSO replicates) stained for H3K9me3, H3K27me3, H3K4me1, H3K27ac. Scatter plots depicting the first 2 discriminant factors derived from features of all four histone modification images for each cell population. Confusion matrix showing classification results of the discriminant analysis. Left column details number of compounds or DMSO replicates for each category in the test set (1 replicate per compound). Numbers represent the percent of compounds classified correctly (diagonal) and incorrectly (off the diagonal). (B) Quadratic discriminant analysis using texture features derived from images of GBM2 cells treated with either DMSO or 85 active compounds (2 technical replicates per compound; 38 DMSO replicates) stained for H3K9me3, H3K27me3, H3K4me1, H3K27ac. Confusion matrix showing classification results from discriminant analysis. Number of compounds or DMSO replicates for each category (1 replicate per compound) is as shown in “b”. Numbers represent the percent of compounds classified correctly (diagonal) and incorrectly (off the diagonal).

**Supplementary Fig. 4.**
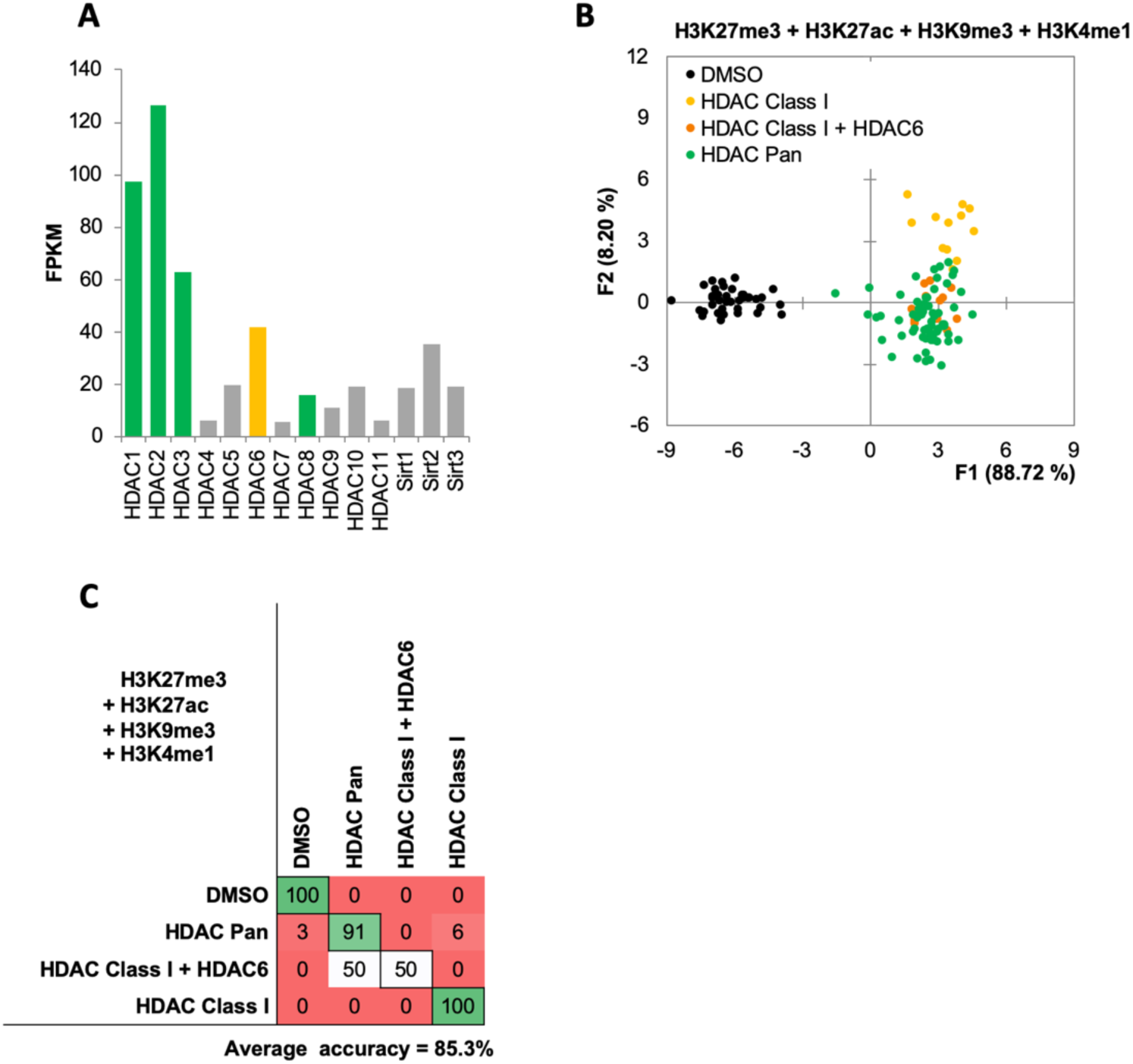
(A) Bar graph showing expression level (FPKM) of HDAC and SIRT genes in GBM2 cells obtained from RNA sequencing. (B, C) Quadratic discriminant analysis using texture features derived from images of GBM2 cells treated with either DMSO or 45 active compounds (2 replicates per compound, 38 DMSO replicates) and stained for H3K9me3, H3K27me3, H3K4me1, H3K27ac. (B) Scatter plot depicting the first 2 discriminant factors derived from features of all histone modification images for each cell population. (C) Confusion matrix showing classification results for the discriminant analysis (test set: 1 replicate per compound; 10 DMSO replicates). Numbers represent the percent of correctly (diagonal) and incorrectly (off the diagonal) classified cell populations.

**Supplementary Fig. 5.**
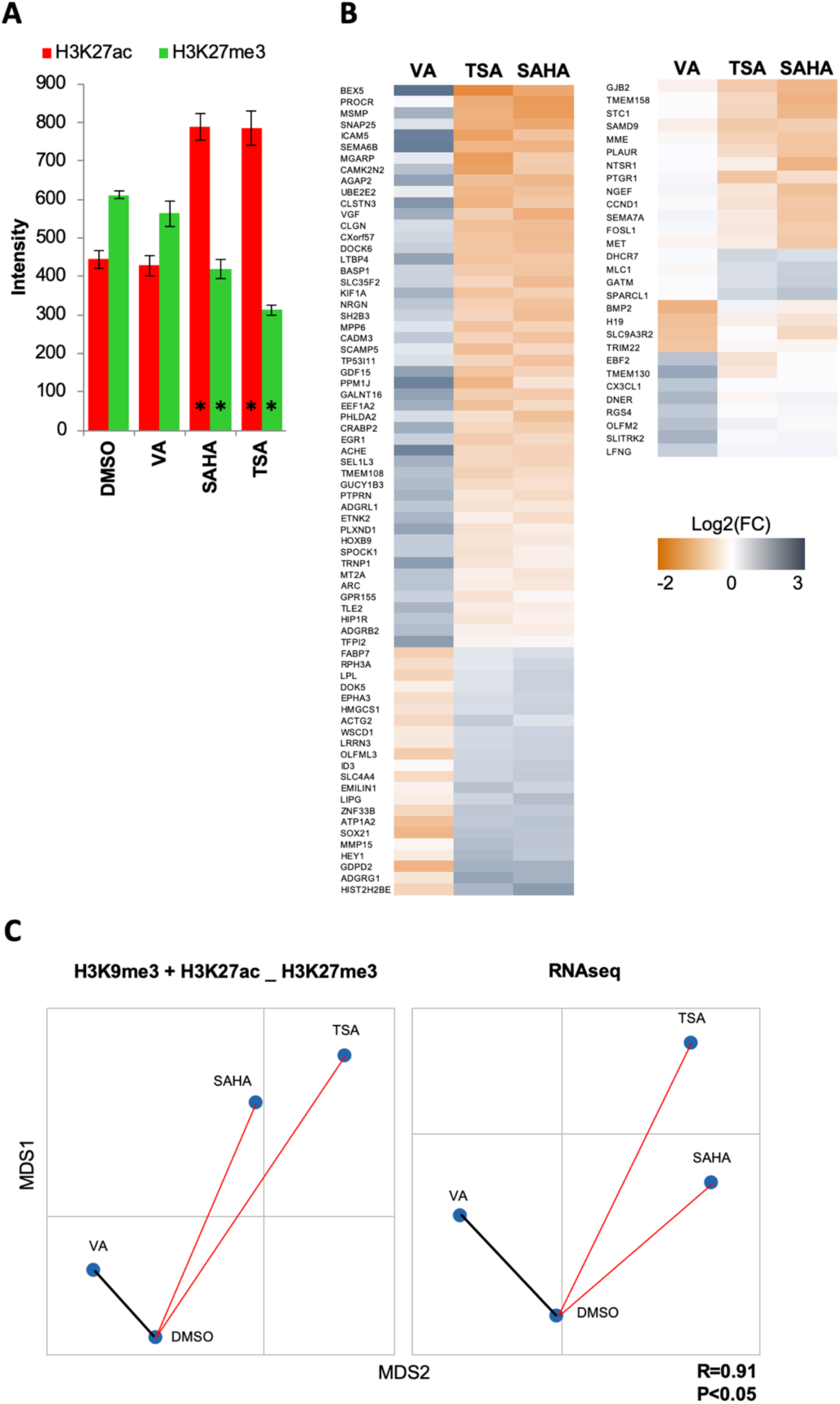
(A) Distance map depicting the relative Euclidean distances between the multiparametric centroids GBM2 cells treated for 24 hours with either DMSO, valproic acid (15 uM), SAHA (3 uM) or TSA (1 uM). Left: Distances calculated using texture features derived from images of H3K9me3, H3K27ac and H3K27me3 marks. Right: Distances calculated using FPKM values of all expressed genes (13,119 genes; FPKM>1 in at least one sample). R denotes Pearson correlation coefficient. (B) Bar graph showing average fold change in average intensity resulting from 24-hour treatment of GBM2 cells with DMSO, valproic acid (15 uM), SAHA (3 uM) or TSA (1 uM) (Mean±SD; n=6 technical replicates). (C) Heat maps showing log2 of fold change in expression (RNA sequencing) of select differentially expressed genes.

**Supplementary Fig. 6.**
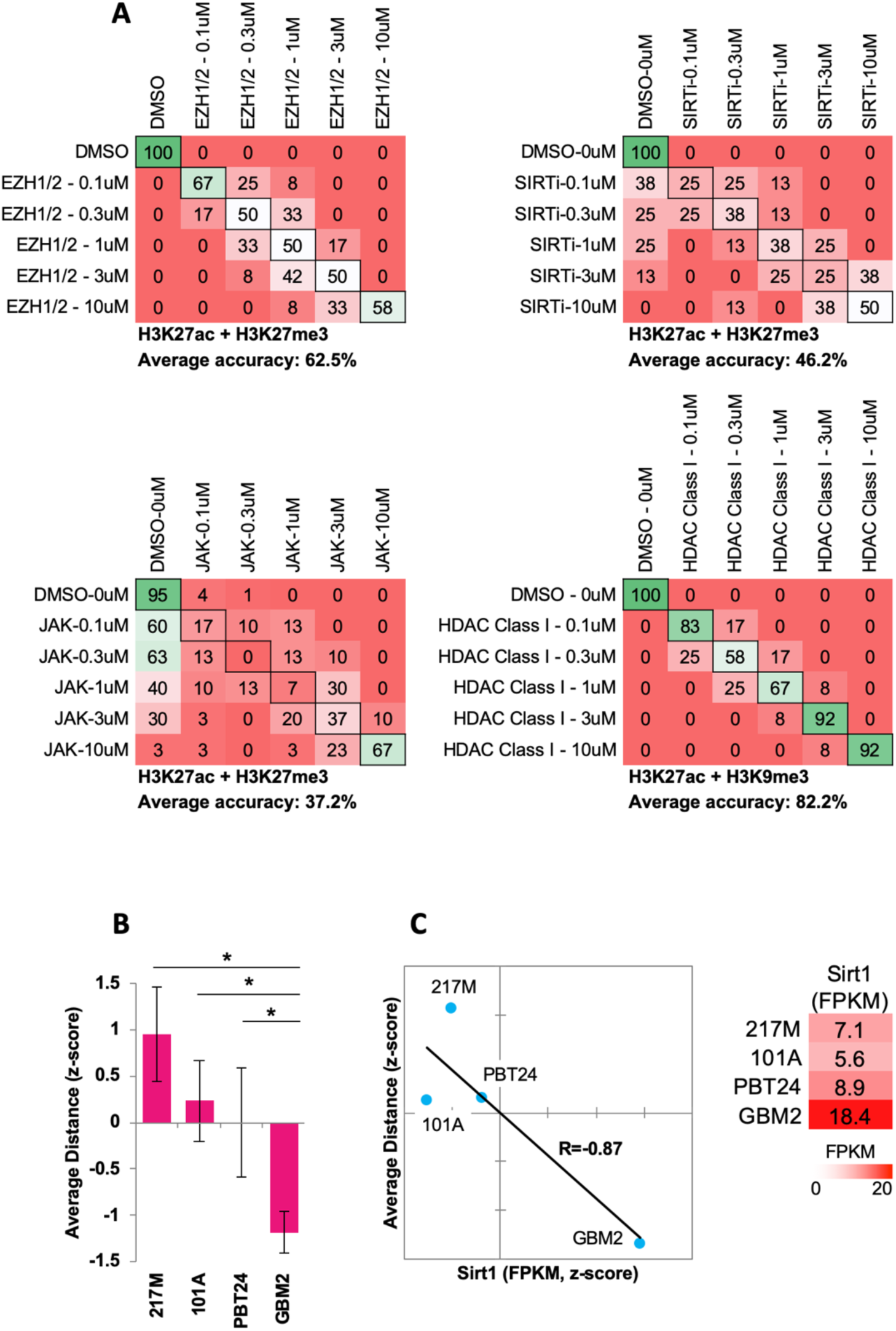
(A) Quadratic discriminant analysis using texture features derived from images of GBM2 cells treated with DMSO, 0.1, 0.3, 1, 3 or 10 uM of EZH1/2 (n=6), SIRT (n=4), JAK (n=15) or HDAC Class I (n=6) inhibitors (2 replicates per compound) stained for either H3K27me3 & H3K27ac (EZH1/2, JAK, SIRT) or H3K9me3 & H3K27ac (HDAC Class I). Confusion matrixes showing results for the discriminant analysis. Numbers represent the percent of replicates classified correctly (diagonal) and incorrectly (off the diagonal). (B) Graph depicting the average z-scored Euclidean distance from DMSO replicates induced by SIRT inhibitors (n=4 compounds, 3 replicates per compound), as calculated using image texture features derived from images of 217M, 101A, PBT24 and GBM2 cells stained for H3K27ac & H3K27me3. (C) Left: Scatter plot comparing the average Euclidean distances shown in “b” with Sirt1 expression in each cell line (z-scored FPKM values derived by RNA sequencing). Right: table showing FPKM values for Sirt1 in the 4 GBM lines. (D) Distance map depicting the relative Euclidean distance between the multiparametric centroids of 4 GBM lines treated with either DMSO, TSA (1 uM), SAHA (3uM) or Tubacin (10 uM). Distances calculated using texture features derived from images of cells stained with H3K9me3 and H3K27ac (n=12 DMSO replicates; n=3 replicates per compound). (e) Polar plot visualizing the fold changes in feature values for cell populations shown in “d” following linear normalization to DMSO averages of each cell line (see Methods).

**Supplementary Fig. 7.**
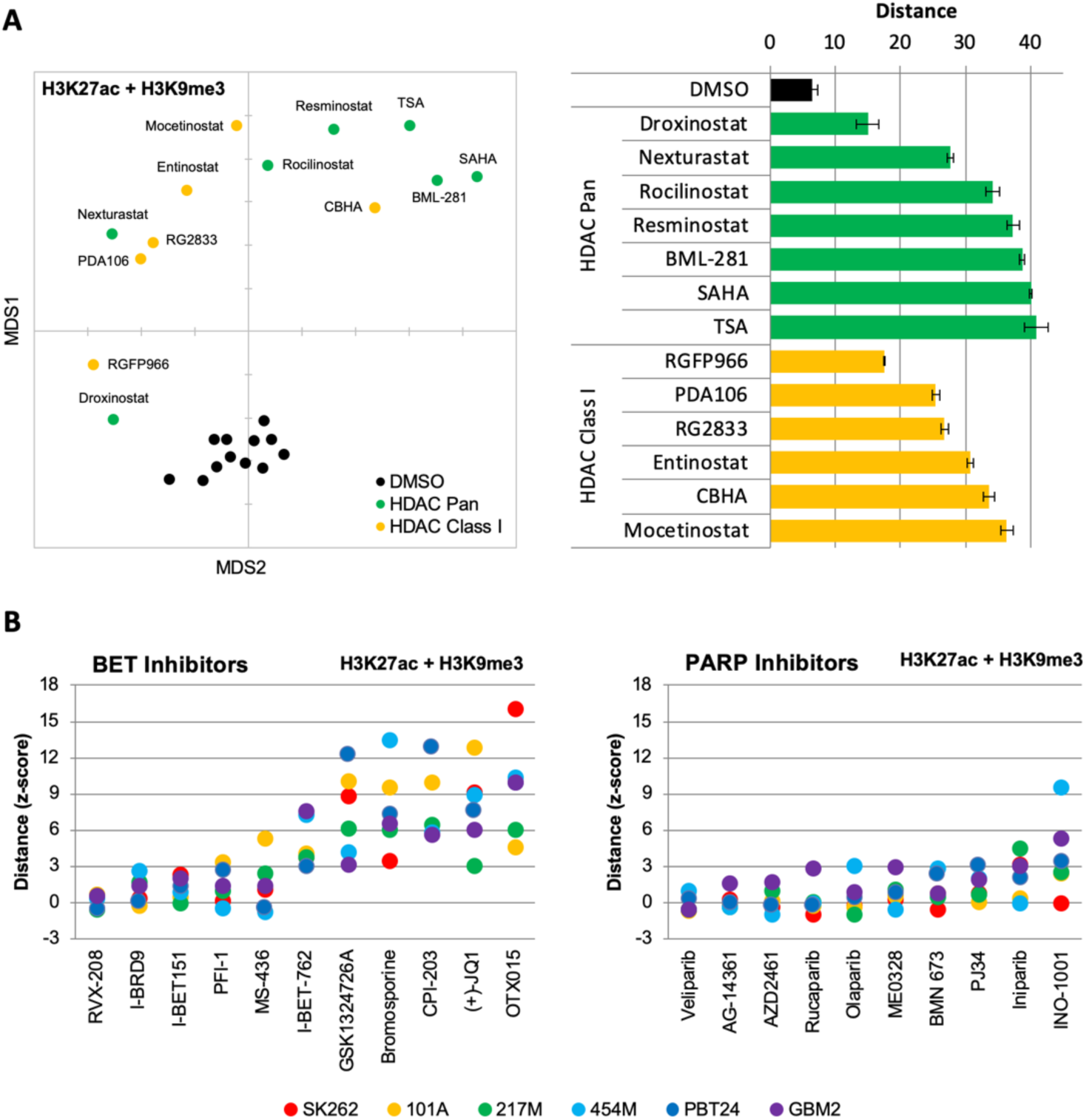
(A) Left: Distance map depicting the relative Euclidean distance between the multiparametric centroids of GBM2 cells treated with either DMSO (n=12 replicates), HDAC Pan (n=7 compounds; showing average or 3 replicates) or HDAC Class I inhibitors (n=6 compounds; showing average or 3 replicates). Distances calculated using texture features derived from images of cells stained with H3K9me3 and H3K27ac. Right: Bar graph depicting the Euclidean distance from DMSO replicates (Mean±SD; n=3 technical replicates) induced by drug treatments shown in “A”. (B) Graph depicting the average fold change in Euclidean distance from DMSO replicates induced by individual BET (left) and PARP (right) inhibitors as calculated using texture features derived from images of H3K27ac & H3K9me3 (n=3 replicates per compound).

**Supplementary Fig. 8.**
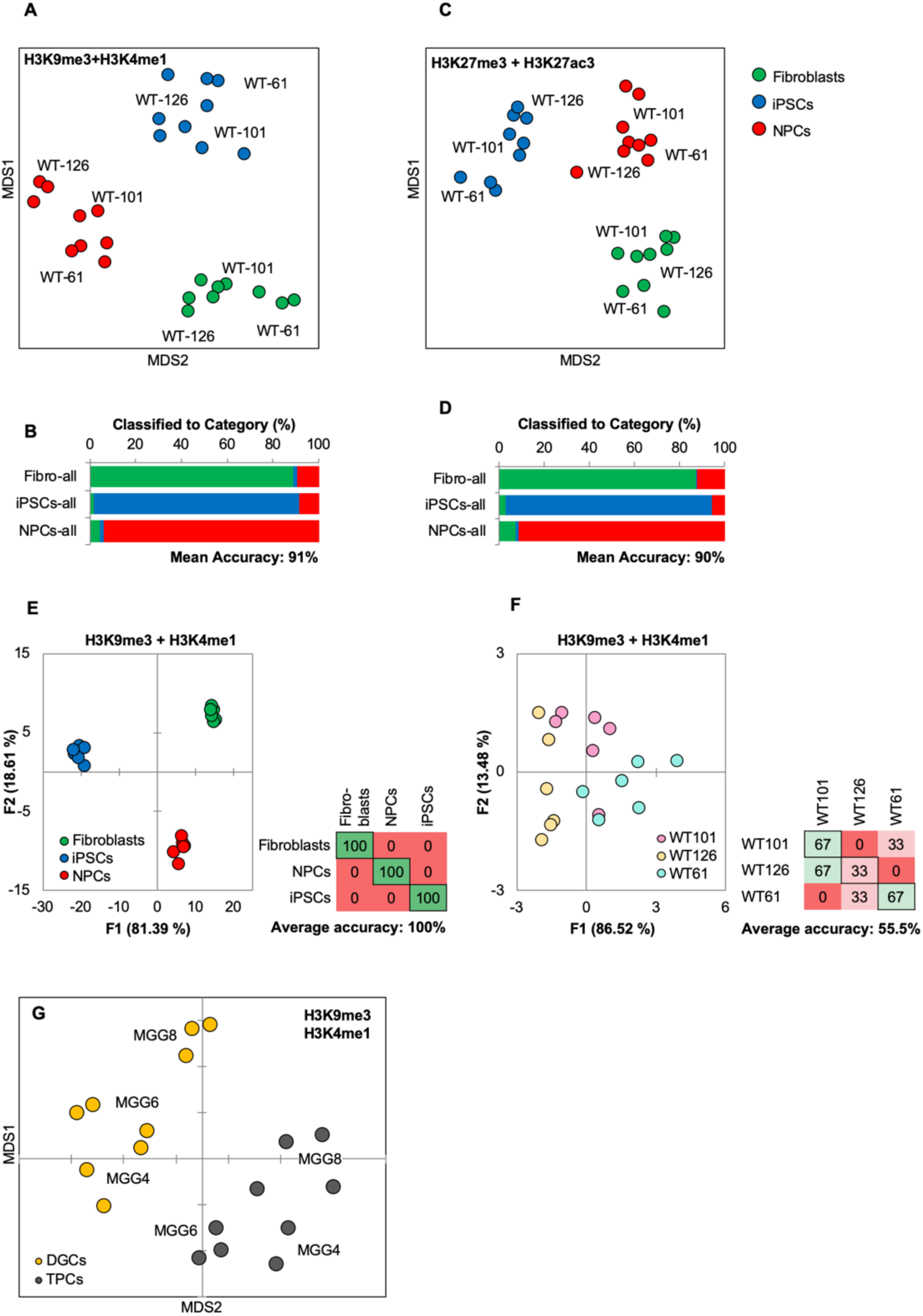
(A, B) Distance map depicting the relative Euclidean distance following MDS between the multiparametric centroids of 9 cell lines: 3 fibroblasts, 3 iPSCs, and 3 NPCs, calculated from texture feature values derived from images of (A) H3K9me3 and H3K4me1 or (B) H3K27ac and H3K27me3 marks. Each cell line appears as 3 technical triplicates. (C, D) Three-way classifications of the 9 cell lines using an SVM classifier trained on image texture features derived from images of pooled fibroblasts, iPSCs, and NPCs stained for (c) H3K9me3 and H3K4me1 or (d) H3K27ac and H3K27me3. (E, F) Quadratic discriminant analysis separating either cell fates or cell lines using texture features derived from images of fibroblasts, iPSCs, and NPC l lines from 3 human donors (WT-61, WT-101 and WT-126; 3 technical replicates each); stained for H3K27me3 and H3K27ac. (E) Discriminant analysis separating the different cell types. Scatter plot depict the first 2 discriminant factors for each cell population (2 replicate per cell line and cell type). Confusion matrixes showing results of classification for the discriminant analysis (test set: 1 replicate per cell line and cell type) Numbers represent the percent of correctly (diagonal) and incorrectly (off the diagonal) classified cell populations. (F) Discriminant analysis attempting to separate the different cell lines. Scatter plot depicting the first 2 discriminant factors for each cell population (2 replicate per cell line and cell type). Confusion matrixes showing results of classification for discriminant analysis (test set: 1 replicate per cell line and cell type). Numbers represent the percent of correctly (diagonal) and incorrectly (off the diagonal) classified cell populations. (G) Distance map depicting the relative Euclidean distance between the multiparametric centroids of 3 genetically distinct TCP and DGC lines calculated using texture features derived from images of H3K9me3 and H3K4me1 marks.

**Supplementary Fig. 9.**
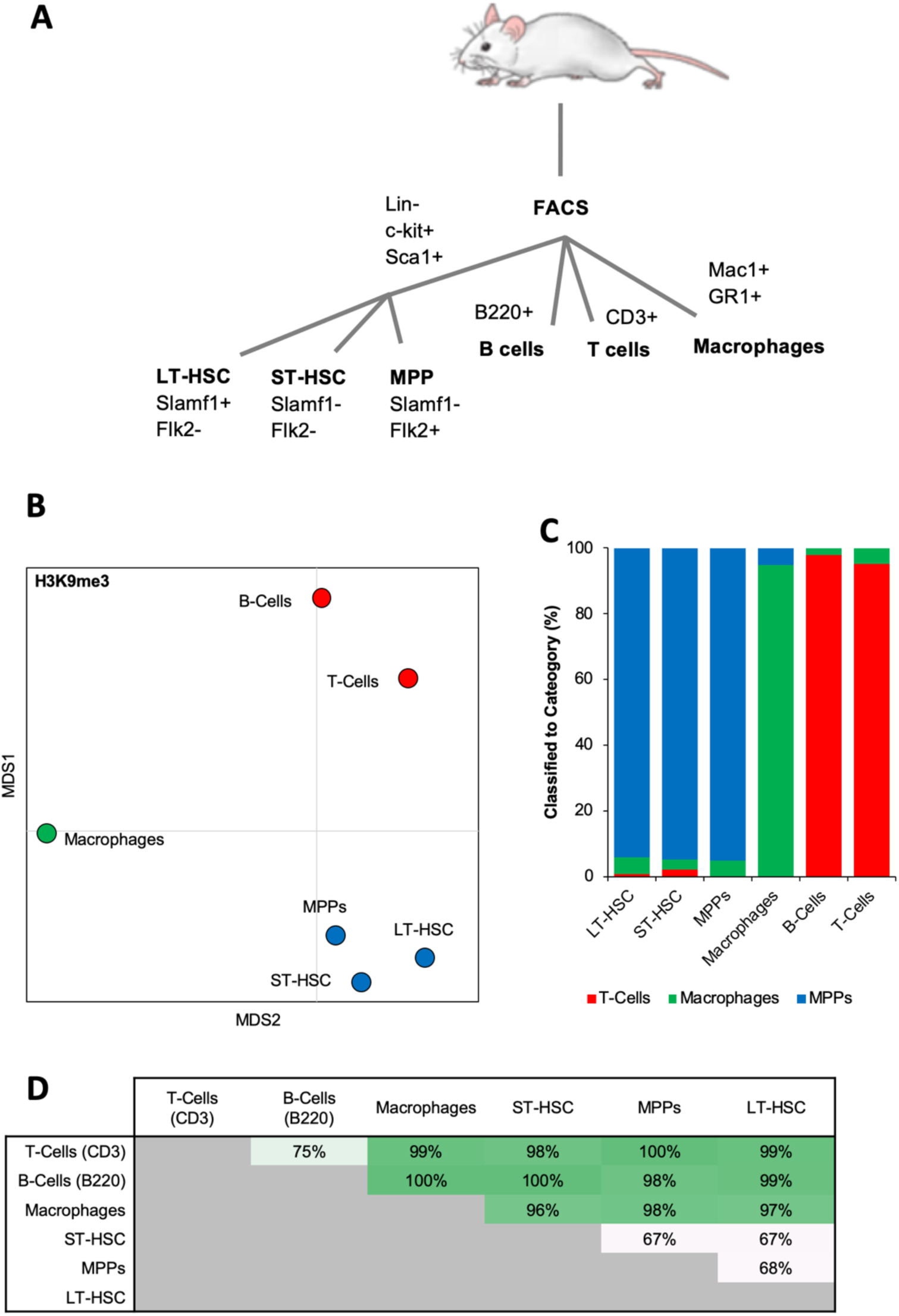
(A) Surface markers for isolation of hematopoietic cells by flow cytometry. (B) Distance map depicting the relative Euclidean distance between the multiparametric centroids of image texture features from immunofluorescence micrographs of 6 hematopoietic cell types. (C) Three-way classification of hematopoietic stem or progenitor cells, T and B lymphoid cells, and macrophages, using an SVM classifier trained on randomly selected subsets of MPPs, macrophages, and T-cells. (D) Accuracy of pairwise SVM classification between the 6 hematopoietic cell types.

**Supplementary Fig. 10.**
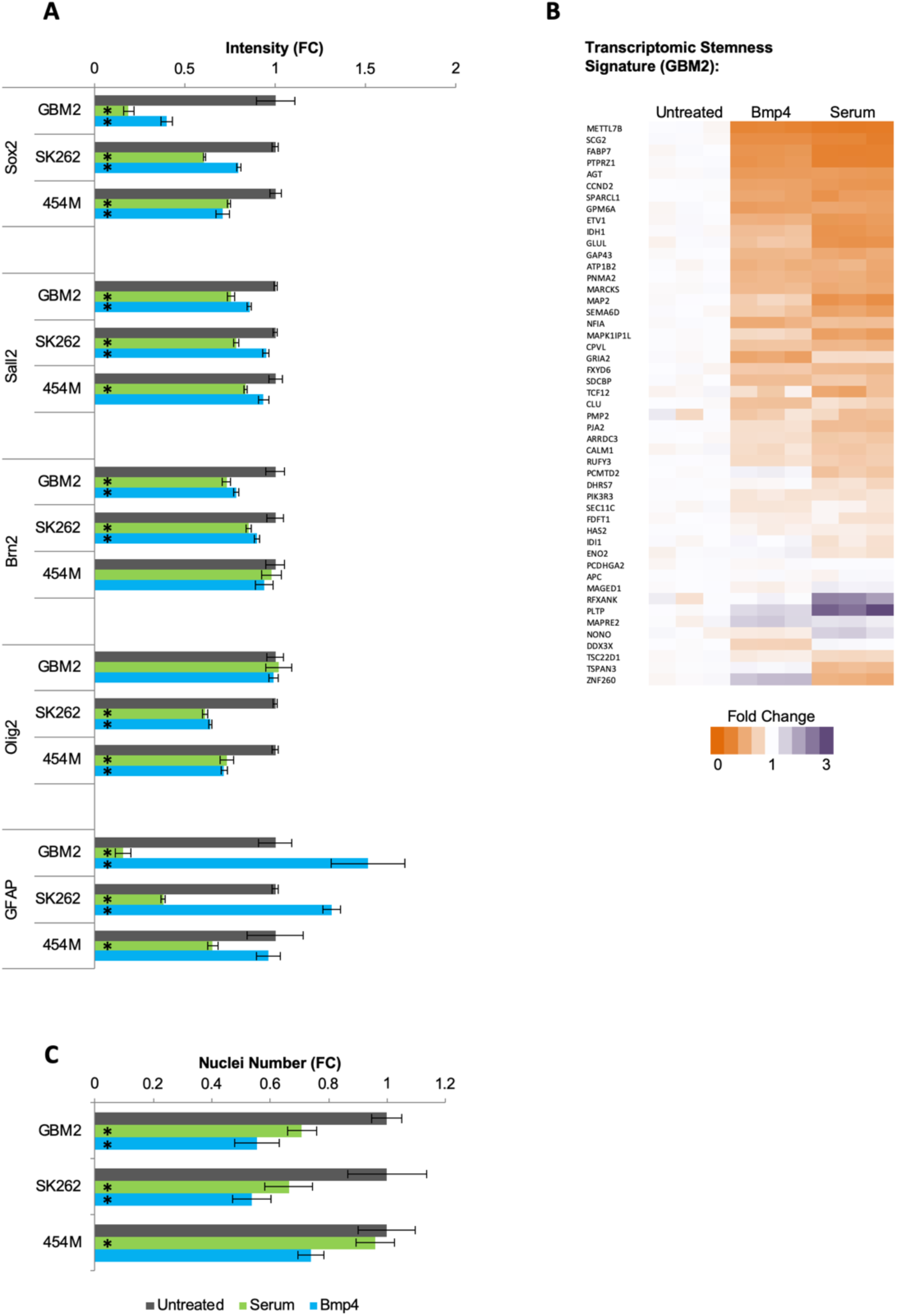
(A) Fold change in immunofluorescence intensity for Sox2, Sall2, Brn2, Olig2, and GFAP for 3-daysserum- or Bmp4-treated primary human GBM lines compared to untreated cells (mean ± S.D, n=3, *p<0.05, unpaired two-tailed t-test). (b) Fold change in nuclei number of 3 days serum- or Bmp4-treated primary human GBM lines compared to untreated cells (mean ± S.D, n=3, *p<0.05, unpaired two-tailed t-test). (c) Heat map showing expression changes of genes identified as the TPC stemness signature in GBM2 cells following 3 days treatment with either serum or Bmp4 (values shown as fold change: FPKM value in every sample divided by average FPKM value of the 3 untreated samples).

**Supplementary Fig. 11.**
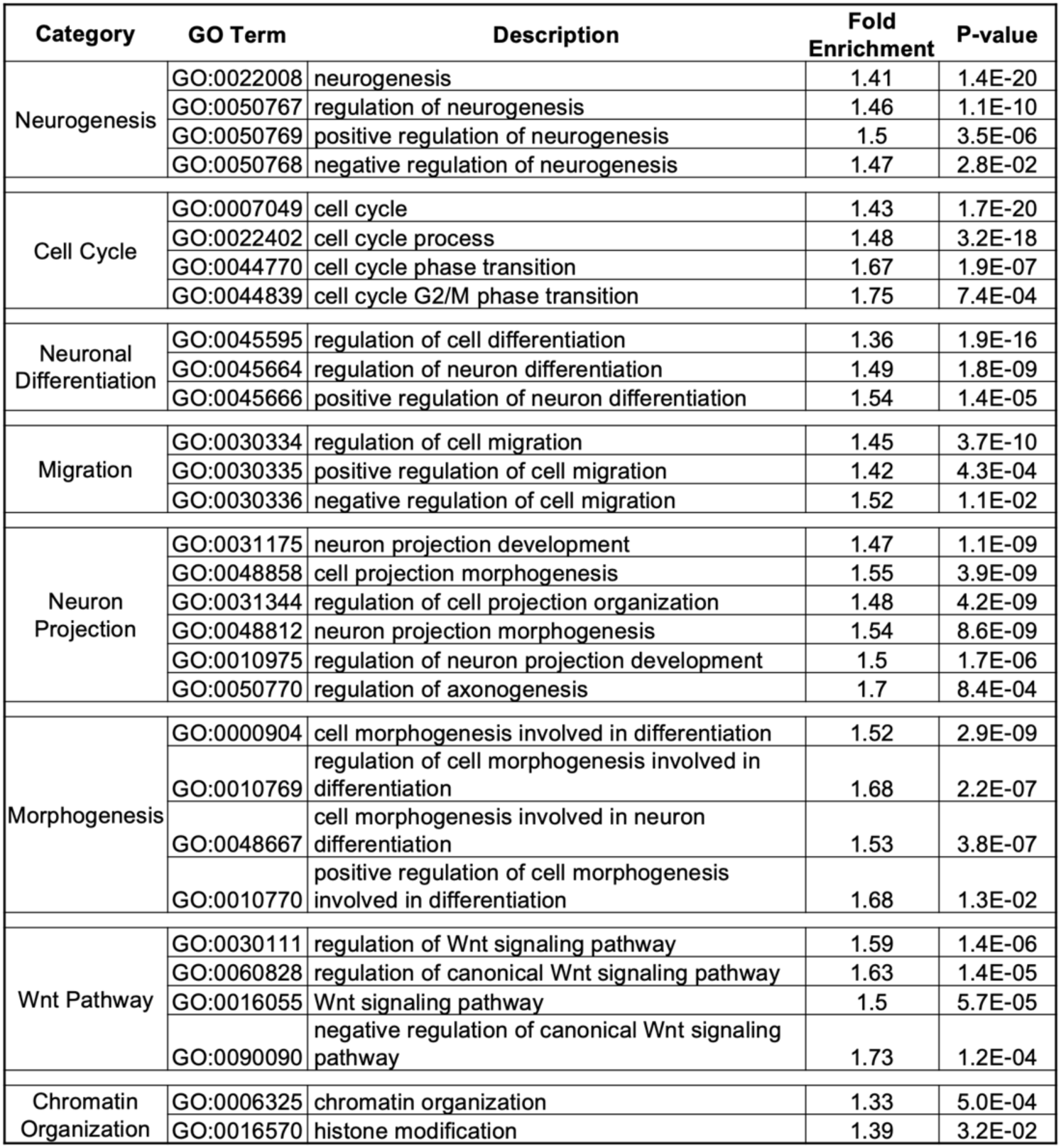
Gene Ontology (GO) terms enriched by serum and Bmp4 treatments and identified using PANTHER v11.

**Supplementary Fig. 12.**
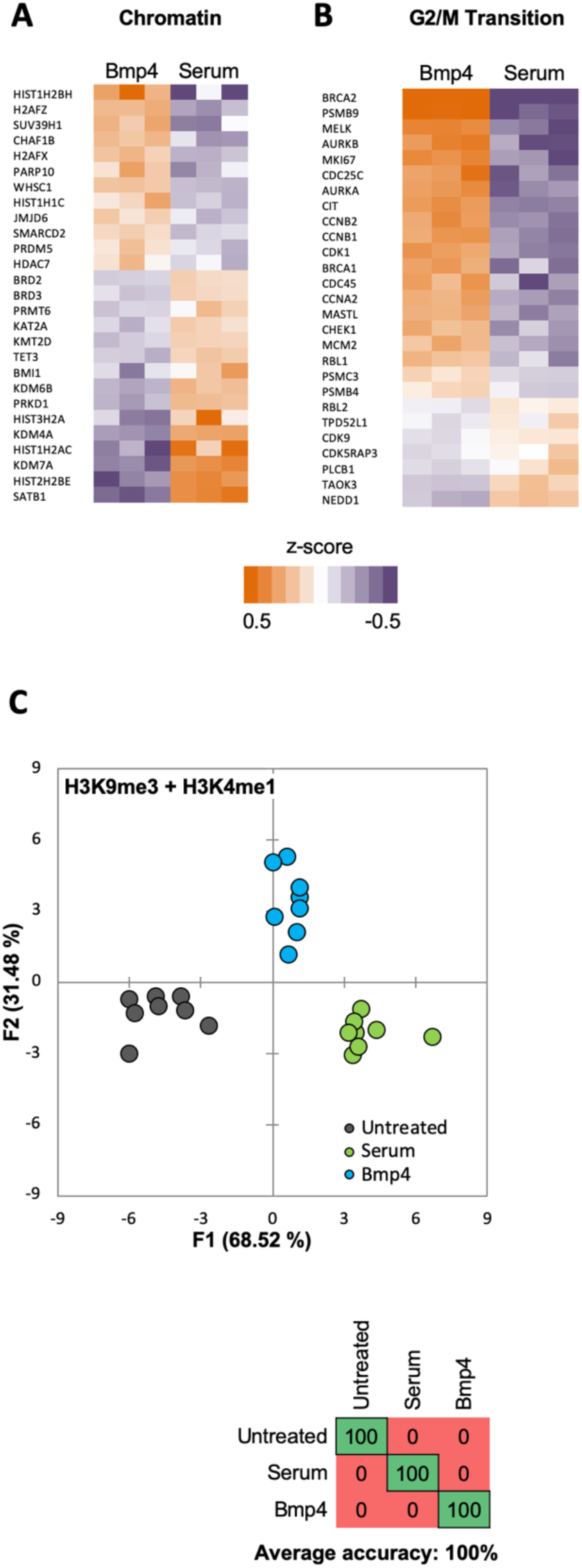
(A, B) Heat maps showing differential expression of selected genes from Gene Ontology (GO) terms (A) chromatin-modification (GO:0006325) or (B) cell-cycle G2/M phase transition (GO:0044839). Expression levels (FPKM) are represented as z-score to highlight difference in levels of expression. (C) Quadratic discriminant analysis using texture features derived from images of untreated or 2 days serum- or Bmp4-treated GBM2, 101A, SK262 and 454M cells (3 replicates per cell lines per treatment) and stained for H3K27me3 and H3K27ac. Scatter plot depicting the first 2 discriminant factors for each cell population (2 replicates per cell lines per treatment). Confusion matrix showing classification results for the discriminant analysis (test set: 1 replicate per cell line per treatment). Numbers represent the percent of correctly (diagonal) and incorrectly (off the diagonal) classified cell populations.

**Supplementary Fig. 13.**
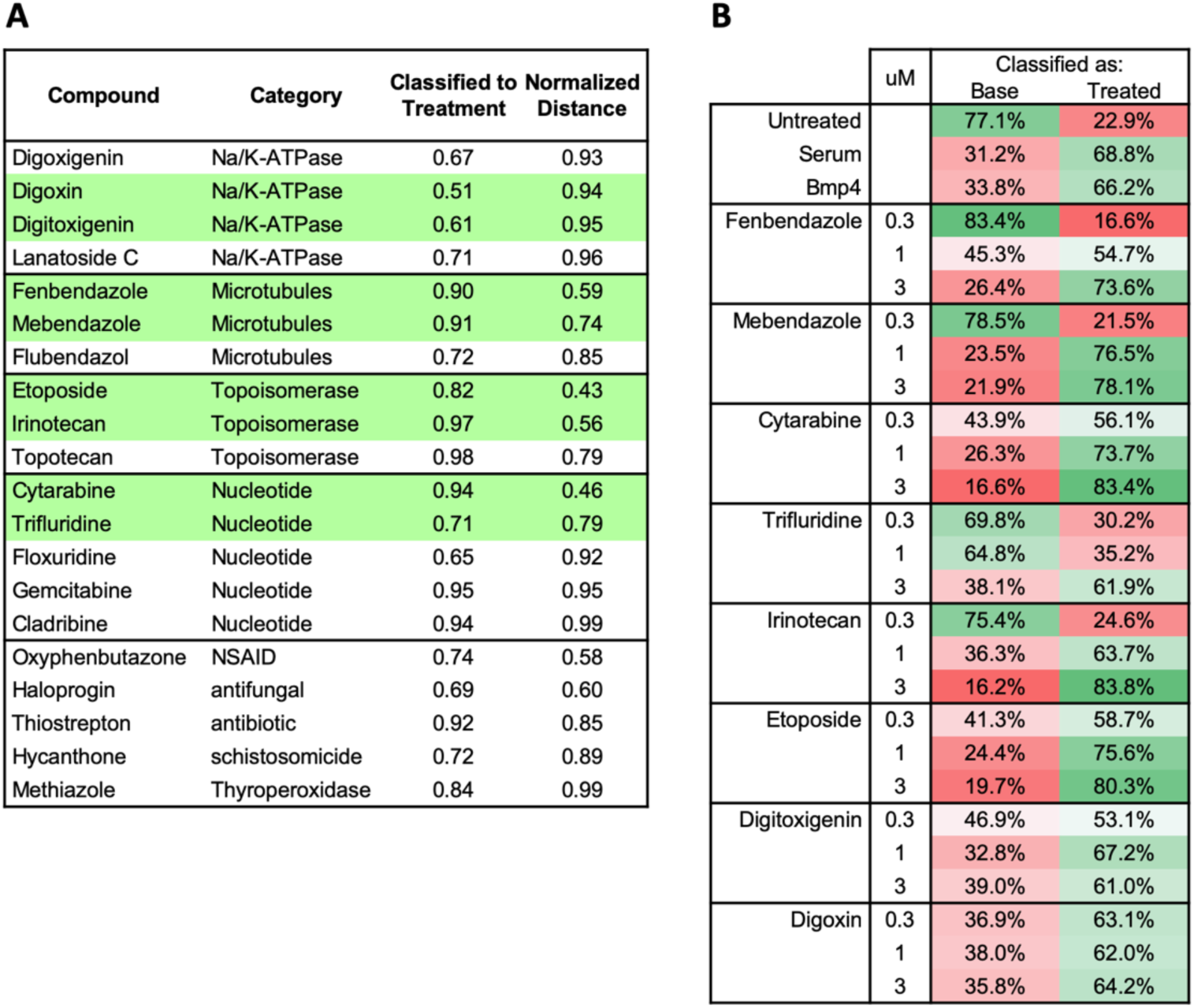
(A) Twenty hit compounds grouped by the functional classes. For the pairwise classification, the classifier was trained on texture features derived from H3K27ac and H3K27me3 images of serum- or Bmp4-treated GBM2 (vs untreated; cut off = classified to treatment>50%). Normalized distance calculated as the Euclidean distance of a compound to either serum or Bmp4 (the smaller of the two) divided by the distance of untreated cells to the same control (cutoff = normalized distance<1). (B) Table showing pairwise classification of indicated drug-treated GBM2 using a classifier trained on texture features derived from H3K27ac and H3K27me3 images of DMSO- and either serum- or Bmp4-treated GBM2 cells.

**Supplementary Figure 14.**
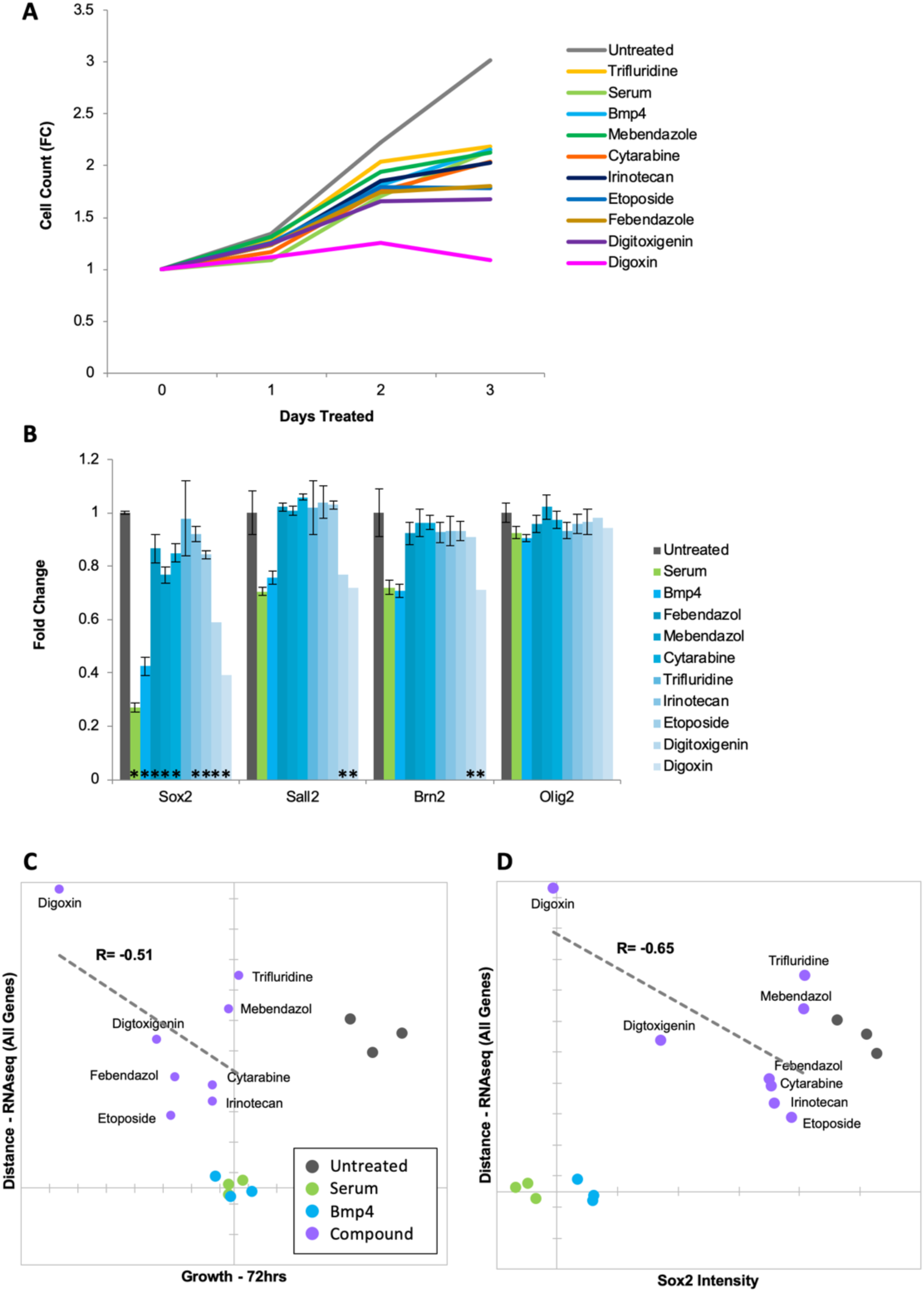
(A) Growth dynamics (fold change in cell count – vertical axis) of untreated, serum-, Bmp4- or drug-treated GBM2 cells over 3 days. (B) Fold change in Sox2, Sall2, Brn2, and Olig2 immunofluorescence intensity of untreated or serum-, Bmp4- or drug-treated GBM2 cells; 3 days of treatment (mean ± S.D, n=3, p<0.05, unpaired two-tailed t-test). (C) Scatter plot showing the correlation of gene expression profile-based ranking and growth rates for untreated, serum-treated, Bmp4-treated, or 8-drugs-treated GBM2 cells. Euclidean distance to serum- or Bmp4-treated GBM2 cells was calculated using transcriptomic profiles (vertical axis), or growth rate after 72 hours treatment with immunofluorescence intensity (horizontal axis). Distances and growth rates were normalized to untreated and serum- or Bmp4-treated GBM2 cells. R denotes Pearson correlation coefficient. (D) Scatter plot showing the correlation of gene expression profile-based ranking and Sox2 expression for 8 candidate drugs, untreated, serum- or Bmp4-treated GBM2 cells. Euclidean distance to serum- or Bmp4-treated GBM2 cells was calculated using transcriptomic profiles (vertical axis), or Sox2 immunofluorescence intensity (horizontal axis). Distances and Sox2 levels were normalized to untreated and serum-Bmp4-treated GBM2 cells.

## Materials and Methods

### Cell Culture

Monolayer cultures of patient-derived GMB TPCs were propagated on Matrigel-coated plates in DMEM:F12 Neurobasal Medium (1:1; Gibco), 1% B27 supplement (Gibco), 10% BIT 9500 (StemCell Technologies), 1 mM glutamine, 20 ng/ml EGF (Chemicon), 20 ng/ml bFGF, 5 µg/ml insulin (Sigma), and 5 mM nicotinamide (Sigma). The medium was replaced every other day and the cells were enzymatically dissociated using Accutase prior to splitting. Fibroblasts, iPSCs, and iPSC-derived NPCs were cultured as previously described (75, 76).

### Differentiation treatment

For TPC differentiation treatments cells were cultured in DMEM:F12 Neurobasal Medium (1:1), 1% B27 supplement, 10% BIT 9500, 1 mM glutamine supplemented with either Bmp4 (100ng/ml; R&D Systems) or FBS (10%).

### Immunofluorescence

Cells were rinsed with PBS and fixed in 4% paraformaldehyde in PBS for 10 min at room temperature. After blocking with PBSAT (2% BSA and 0.5% Triton X-100 in PBS) for 1 hour at room temperature, the cells were incubated overnight at 4°C with primary antibodies diluted in PBSAT. Primary antibodies are listed in Table 1, and the appropriate fluorochrome-conjugated secondary antibodies were used at 1:500 dilution. Nuclear co-staining was performed by incubating cells with either Hoechst-33342 or DAPI nuclear dyes.

### Microscopy and image analysis

For MIEL analysis, cells were imaged on either an Opera QEHS high-content screening system (PerkinElmer) using ×40 water immersion objectives or an IC200-KIC (Vala Sciences) using a ×20 objective. Images collected were analyzed using Acapella 2.6 (PerkinElmer). At least 40 fields/well for Opera and 5 fields/well for IC200 were acquired and at least 2 wells per population were used. Features of nuclear morphology, fluorescence intensity inter-channel co-localization, and texture features (Image moments, Haralick, Threshold Adjacency Statistics) were calculated using custom algorithms (scripts available from www.andrewslab.ca). A full list of the features used is available from the authors. Values for each cell were generated and exported to Microsoft Excel or MATLAB for further analysis. For Sall2, Olig2, Brn2, Sox2, Oct4, and GFAP immunostaining, images were captured on an IC200-KIC (Vala Sciences) using a ×20 objective. Between 3 and 8 fields per well were acquired and analyzed using Acapella 2.6 (PerkinElmer). For all nuclear markers, average intensities in nucleus or fold change compared to untreated cells are shown. Unless stated otherwise, at least 3 wells and a minimum of 300 cells for each condition were compared using the unpaired two-tailed t-test.

### Data processing

The image features-based profile for each cell population (e.g., cell types, treatments, technical repetition) was represented using a vector (center of distribution vectors) in which every element is the average value of all cells in that population for a particular feature. The vector’s length is given by the number of features chosen (262 per histone modification). Raw feature values were normalized by z-scoring to the average and standard deviation of all populations being compared. All cells in each population were used to calculate center vectors, and each population contained at least 50 cells. Activity level for each drug was determined by calculating the distance from DMSO. For this, feature values of all DMSO replicates center vectors were used to calculate the DMSO center vector. Euclidean distance of each compound and each DMSO replicate to the DMSO center vector was calculated. Distances were z-scored to the average distance and standard deviation of DMSO replicates from the DMSO center vector. Transcriptomic-based profile for each cell population was represented using a vector in which every element is the z-scored FPKM value for a single gene in that population. The length of the vector is given by the number of genes used to construct the profile.

### Multidimensional scaling - MDS

The Euclidean distance between all vectors (either image features or transcriptomic based) was calculated to assemble a dissimilarity matrix (size N×N, where N is the number of populations being compared). For representation, the N×N matrix was reduced to a Nx2 matrix with MDS using the Excel add-on program Xlstat (Base, v19.06), and displayed as a 2D scatter plot.

### Discriminant Analysis

Quadratic discriminant analysis was conducted using the Excel add-on program xlstat (Base, v19.06). The model was generated in a stepwise (forward) approach using default parameters. All features derived from images of tested histone modification were used for analysis following normalization by z-score. Features displaying multicollinearity were reduced. Model training was done using multiple DMSO replicates and at least 2 replicates from each cell-line or drug treatment. The model was tested on at least 8 DMSO replicates and at least 1 replicate from each cell line or treatment.

### SVM classification

SVM classification was conducted as previously described (30). Cell-level data in total populations (minimum 400 cells per population) were normalized to z-scores, and a subset of cells from each population being classified was randomly chosen as the training set (subset size at least 100× the population number bei ng classified). The training set was used for a SVM classifier (MATLAB svmtrain function). The remaining cells (test set) were then classified using the SVM-derived classifier to assess the accuracy of classification (MATLAB svmclassify function). Here, the accuracy of all pairwise classifications was given as the average accuracy calculated for each population. To classify the similarity of multiple cell populations, we classified known populations (e.g., different treatments or cell fates) to generate known bins and then used the same classifiers on the unknown population to categorize each cell.

### Epigenetic Drug Screening

GBM2 cells were plated at 4000 cells/well and exposed to epigenetic compounds (Table 2) at 10 µM for 1 day in 384-well optical bottom assay plates (PerkinElmer). Negative control was DMSO (0.1%), 48 DMSO replicates per plate, 3 technical replicates (wells) were treated per compound. Cells were fixed and stained with histone modification-specific antibodies (H3K27ac & H3K27me3, H3K9me3, H3K4me1) and AlexaFluor-488- or AlexaFluor-555-conjugated secondary antibodies. DNA was stained with DAPI followed by imaging and feature extraction. To compare data from multiple plates, average feature values in each plate were normalized to DMSO. Here, feature values of all DMSO replicates center vectors in each plate, then were used to calculate the plate-wise DMSO vector. Raw feature values for all center vectors of all populations in each plate were normalized to the plate-wise DMSO vector; normalized feature values were z-scored as above. To identify active compounds, activity level for each compound was calculated as above, and active compounds were defined as compounds for which activity z-score was >3. Compounds reducing the number of imaged cells per well below 50 were considered toxic and excluded from analysis.

### Concentration Curves

GBM2 cells were plated and stained as above. For each compound (Table 3), cells were treated at 0.1, 0.3, 1.0, 3.0, 10.0 uM. Activity levels were calculated as above. Average cell count was calculated across the replicates for each compound to compare epigenetic changes and toxicity. Cell counts were z-scored against the average and standard deviation of all DMSO replicates. Distances (z-scored) and cell counts (z-scored) were averaged for each functional class at each concentration.

### RNAseq and transcriptomic analysis

Total RNA was isolated from GBM2 cells using the RNeasy Kit (Qiagen), 0.25 ug total RNA was used to isolate mRNAs and for library preparation. Library preparation and sequencing were conducted by the SBP genomics core (Sanford-Burnham NCI Cancer Center Support Grant P30 CA030199). PolyA RNA was isolated using the NEBNext® Poly(A) mRNA Magnetic Isolation Module, and barcoded libraries were made using the NEBNext® Ultra II™ Directional RNA Library Prep Kit for Illumina®(NEB, Ipswich MA). Libraries were pooled and single-end sequenced (1X75) on the Illumina NextSeq 500 using the High-Output V2 kit (Illumina). Read data, processed in BaseSpace (basespace.illumina.com), were aligned to Homo sapiens genome (hg19) using STAR aligner (https://code.google.com/p/rna-star/) with default settings. Differential transcript expression was determined using the Cufflinks Cuffdiff package (https://github.com/cole-trapnell-lab/cufflinks). For heat maps showing fold change in expression, FPKM values in each HDACi-treated population were divided by the average FPKM values of DMSO-treated GBM2 and values shown as log2 of the ratio. Go enrichment analysis was conducted using PANTHER v11 (77) using all genes identified as differentially expressed following either serum or Bmp4 treatment. To highlight differences in expression levels between serum- and Bmp4-treated GBM2 cells, FPKM values in each sample were z-scored. Zscore=(FPKM_Observation_-FPKM_Average_)/FPKM_SD_ (FPKM_Observation_-FPKM value obtain through sequencing; FPKM_Average_ – average of all FPKM values in all samples for a certain gene; FPKM_SD_ – standard deviation of FPKM values for a certain gene). Heat maps were generated using Microsoft Excel conditional formatting.

### Comparing epigenetic changes in different cell lines

To compare drug-induced epigenetic changes across multiple glioblastoma cell lines, 101A, 217M, GBM2 and PBT24 cells were plated at 4000 cells/well and treated with compounds for 24 hours. Compounds and concentrations are shown in Table 4. Activity level was calculated as above. Pearson coefficient and significance of correlation for activity levels in each pair of cell lines were calculated using the Excel add-on program xlstat (Base, v19.06).

### Correlation of transcriptomic and image-based profiles

Euclidean distances were calculated using either transcriptomic data (FPKM) or texture features. Pearson’s correlation coefficient (R) was transformed to a t-value using the formula (t = R × SQRT(N-2)/SQRT(1-R2) where N is the number of samples, R is Pearson correlation coefficient; the p-value was calculated using Excel t.dist.2t(t) function. For compound prioritization, Euclidean distance between the compound treated and serum- or Bmp4-treated GBM2 cells was calculated based on either FPKM)or image features. The average distance for both serum and Bmp4 treatments was normalized to the average distance of untreated cells to serum and Bmp4.

### Sensitization to radiation or TMZ

Cells were plated at 1500 cells/well in 384-well optical bottom assay plates (PerkinElmer). Two sets of the experiment were prepared; DMSO (0.1%) was used for negative controls at 48 DMSO replicates per plate; 3 replicates (wells) were treated per compound. Compound concentrations used are shown in Table 5. Cells in both sets were pre-treated with epigenetic compounds for 2 days prior to cytotoxic treatment. Cytotoxic treatment, either 200uM temozolomide (TMZ, Sigma) or 1Gy x-ray radiation (RS2000; RAD Source) was carried out for 4 days on single set (‘treatment set’); for TMZ treatment, DMSO control was given to the second set. A single radiation dose was given at day 3; TMZ was given twice at days 3 and 5 of the experiment. Cells were fixed, stained with DAPI, and scored using an automated microscope (Celigo; Nexcelom Bioscience). For each compound, fold change in cell number was calculated for both the “treatment set” (Drug+Cytotox) and the “control set” (Drug), compared to DMSO-treated wells in the control set. The effect of radiation or TMZ alone was calculated as fold reduction of DMSO-treated wells in the treatment set compared to DMSO-treated wells in the control set (Cytotox). The coefficient of drug interaction (CDI) was calculated as (Drug+Cytotox)/ (Drug)X(Cytotox). For conformation experiments, the same regiment and CDI calculations were carried out on SK262, 101A, 217M, 454M, and PBT24 glioblastoma cell lines; PARPi and BETi were used at same concentration as the initial screen on GBM2 (Table 5).

### Prestwick Chemical Library screen using H3K27me3 and H3K27ac

GBM2 cells were plated at 2000 cells/well and exposed to Prestwick compounds (3 µM; Table 6) for 3 days in 384-well optical bottom assay plates (PerkinElmer). Cells were then fixed and stained with rabbit polyclonal anti-H3K27ac and mouse monoclonal anti-H3K27me3 antibodies followed by AlexaFluor-488- or AlexaFluor-555-conjugated secondary antibodies. Positive controls contained BMP4 (100 ng/ml) and serum (10%); negative controls contained DMSO (0.1%). DNA was counterstained with Hoechst. Images were acquired using Perkin Elmer Opera® QEHS. MIEL analysis was conducted as described above.

